# GM-CSF modulates conserved ILC-like transcriptional states and myeloid cell signaling during ulceration in Crohn’s disease

**DOI:** 10.64898/2026.02.03.703526

**Authors:** Joshua K. Morrison, Ksenija Sabic, Neha Maskey, Sayali Talware, Nai-yun Hsu, Colleen Chasteau, Elizabeth Aslinger, Jake Herb, Shikha Nayar, Rachel Levantovsky, Christopher Tastad, Rachel Moss, Alan Soto, Monica Garcia-barros, Jessy Ntunzwenimana, Mariel Glass, Michelle Bao, Jiayu Zhang, Huajun Han, Jane Stevens, Lorena Tavares, Tin Htwe Thin, Sergey Khaitov, Alexander Greenstein, Rachel Brody, Jaime Chu, Arthur Mortha, Judy H. Cho, Ling-shiang Chuang

## Abstract

Macrophage (M-), granulocyte (G-), and granulocyte–macrophage (GM-) colony-stimulating factors (CSFs) regulate myeloid cell function, yet their relative roles during inflammation remain poorly defined. To investigate how CSFs are spatially organized in Crohn’s disease, we performed Xenium single-cell spatial transcriptomics on ileal tissues, revealing cell-type–specific expression and source–target interactions for each CSF. GM-CSF, unlike M-CSF or G-CSF, was locally enriched in ulcerated regions and associated with STAT5 phosphorylation within macrophage aggregates. To study functional consequences of GM-CSF signaling, we developed a csf2rb⁻/⁻ zebrafish model of intestinal injury. Loss of GM-CSF signaling exacerbated epithelial damage and inflammation, whereas recombinant human GM-CSF limited injury and was associated with a reduced DSS-induced increase in putative ILC1-like cells, preservation of putative ILC3-like cells, and increased IL-22 production. Cross-species single-cell analysis revealed conserved ILC-like transcriptional states and GM-CSF-responsive myeloid transcriptional programs. Together, these findings demonstrate a protective role for GM-CSF during intestinal inflammation and support a model in which spatially localized GM-CSF signaling and associated myeloid and ILC-like cell states contribute to intestinal immune homeostasis in Crohn’s disease.

## Introduction

Colony stimulating factor (CSF) cytokines, including macrophage-CSF (M-CSF/*CSF1*), granulocyte-macrophage-CSF (GM-CSF/*CSF2*), and granulocyte-CSF (G-CSF/*CSF3*) are central regulators of myelopoiesis. While M-CSF and G-CSF are essential for macrophage and neutrophil regulation, respectively, GM-CSF is relatively dispensable during steady-state, apart from supporting alveolar macrophages and tissue dendritic cells^1^. During intestinal inflammation, however, GM-CSF promotes myeloid activation, cell migration, and macrophage renewal, making it an important component of the inflammatory response in Crohn’s disease (CD). CD, a chronic inflammatory bowel disease, involves dysregulated myeloid activity^2^, underscoring the importance of dissecting the distinct roles of CSF cytokines in inflammation.

Genome-wide association studies have linked *CSF2RB*, the GM-CSF receptor beta chain, to IBD risk^2,3^. Further, the presence of neutralizing anti-GM-CSF autoantibodies has been associated with increased CD complications^4^. Recent studies suggest GM-CSF, through phosphorylation of STAT5^5^, can induce regulatory states in monocytes and dendritic cells that contribute to epithelial homeostasis^5,6^. However, conflicting pre-clinical results^7–9^, including mice deficient in GM-CSF or CSF2RB and anti-GM-CSF neutralizing antibody administration, highlight the need for a better understanding of its context-dependent functions and cellular targets in IBD. Within the intestine, innate lymphocytes, particularly group 3 innate lymphoid cells (ILC3s), are major GM-CSF producers^6,10^. ILCs orchestrate mucosal immunity through integration of cytokine, stress, and microbial signals, and ILC3s support intestinal epithelial homeostasis via IL-17 and IL-22 production^11^. Expansion of pro-inflammatory, IFN-gamma-producing ILC1s and contraction of ILC3s have been reported during CD inflammation. Furthermore, anti-GM-CSF autoantibody levels correlate with this altered ILC balance^4,12^. These observations suggest an association between GM-CSF signaling and intestinal ILC states, although the cellular mechanisms linking these processes remain unclear.

Many genes associated with CD, particularly those with the largest effect sizes, are expressed by myeloid cells and reflect a genetic architecture shaped by host responses to mycobacterial species^2^. Importantly, the zebrafish pathogen, *Mycobacterium marinum*, has provided substantial *in vivo* insight into human mycobacterial pathogenesis^13,14^. To investigate intestinal inflammation, we employed the dextran sodium sulfate (DSS)-induced intestinal injury model in larval zebrafish—a well-established system for studying epithelial damage and innate immune activation. This model leverages the organism’s optical transparency, conserved immune cell lineages and signaling pathways, and, critically, its lack of functional adaptive immunity at the larval stage, enabling assessment of innate immune cell-driven processes during mucosal injury and repair^15,16^. Moreover, the identification of zebrafish ILC-like cells with transcriptional features analogous to mammalian ILCs provides an experimental framework for investigating GM-CSF-associated innate immune responses^17^.

While single-cell RNA sequencing has advanced our understanding of cytokine signaling in IBD^16,18,19^, it lacks the spatial resolution necessary to contextualize cellular interactions. Given the limited detection of GM-CSF in serum, even during inflammation, and its production by tissue-resident cells^4,6^, understanding local GM-CSF signaling requires integration of single-cell and spatial approaches. We therefore leveraged Xenium single-cell spatial transcriptomics to characterize the cellular sources, potential targets, and spatial organization of CSF signaling within intestinal tissues^20^. Defective GM-CSF signaling, whether caused by neutralizing anti– GM-CSF autoantibodies^4^ or loss-of-function mutations in *CSF2RB*^5^, has been linked to increased susceptibility to CD and greater risk of disease complications. Defining the cell-type specific and spatial context of GM-CSF signaling during intestinal inflammation is therefore important. By integrating Xenium spatial transcriptomics from paired inflamed and uninvolved CD tissues with functional analyses in a *csf2rb*^⁻^***^/^***^⁻^ zebrafish model, we investigated how GM-CSF signaling is associated with myeloid responses and ILC-like transcriptional states during intestinal inflammation.

## Methods

### Ileal tissue collection and processing

Ileocolic resection samples were examined by a pathologist, with transmural sections taken from inflamed, uninflamed, and stricture regions within an hour. Tissues were fixed in 10% formalin for 48 hours, preserved in 70% ethanol for up to 72 hours, and then embedded in paraffin at Mount Sinai’s Biorepository and Pathology Core.

### Xenium In Situ spatial transcriptomics

FFPE blocks were sectioned (5µm) at Mount Sinai’s Biorepository and Pathology Core per 10x Genomics protocol CG000578. Sections were placed on Xenium slides, dried, deparaffinized, and permeabilized. mRNAs were targeted using specific probes, hybridized overnight at 50°C. After washing, probes were ligated (37°C, 2h), then amplified enzymatically to enhance signal. Background fluorescence was quenched, and sections were imaged using the Xenium Analyzer, which performed automated imaging, liquid handling, and analysis. Z-stacks were acquired with a 0.75 μm step size.

### Xenium data processing

Cells were filtered and gene expression data normalized according to Xenium standard preprocessing procedures using the scanpy python package and python version 3.10.4. Filtering parameters included (a) the minimum number of genes expressed per cell and transcript counts per cell (for cell filtering) and (b) the minimum number of transcripts for each gene and number of cells in which genes were expressed (for gene filtering, though no genes were removed as all passed the filtering parameters). The gene expression matrix was total-count-normalized (target sum of 10,000 per cell) and log-normalized. Data were zero-centered and scaled to unit variance for dimensionality reduction and clustering. Additionally, the scaled data were clipped to a maximum value of 10. Each sample was clustered separately using scanpy by performing (a) dimensionality reduction via PCA, (b) UMAP embedding, then (c) Leiden clustering on the scaled gene expression data. Only genes included in the panel as cell type markers were used for clustering. Clustering resolutions of 0.5 (with a minimum distance of 0.5), 0.75 (with a minimum distance of 0.3), and 1.5 (with a minimum distance of 0) were attempted to find the optimal resolution. A resolution of 1.5 was used for analysis. For Xenium ILC analysis, we first calculated the densities (cells/μm²) of ILC1s (IL7R⁺TBX21⁺) and ILC3s (IL7R⁺RORC⁺) from the Xenium dataset. The relative ILC ratio was then determined by calculating the proportion of each ILC subset relative to the total IL7R⁺ ILC population identified in each sample following single-cell clustering and cell-type annotation. Finally, these proportions were normalized to the control condition to enable comparison across treatment groups.

### Ligand-receptor analysis

Ligand-receptor analysis was performed using stlearn (v1.1.5)^21^ in Python 3.10.18 on mucosal single-cell data. Xenium data were gridded into 8×8 micron spots. Based on connectomeDB2020^22^, 22 ligand-receptor pairs were identified. Co-expression was analyzed within a diffusion distance of 150 microns. A background distribution of 20,000 random gene pairs was used for significance testing. P-values were adjusted via Benjamini-Hochberg correction (FDR < 0.05). Ligand-receptor pairs required expression in at least two spots for testing.

### CRISPR-Cas9 mutagenesis in zebrafish

58 pmol CRISPR RNA (crRNA) (5’-CATTGGGTAAAACTACATG-3’), 58 pmol FP-labelled trans-crRNA (Sigma), and 6.75 pmol Cas9 protein (Sigma) were combined and allowed to complex for 30 min on ice before injection. Wild-type AB zebrafish embryos were injected with 4 nl of injection mix per embryo at the 1–4-cell stage. Embryos were screened for the presence of fluorescent guide RNA at 2 hr after fertilization. At 5 days post-fertilization (dpf), larvae were collected for gDNA extraction. A region of genomic DNA containing the intended CRISPR target sites was amplified by PCR (F primer: 5’-CTGCAGTGATGGATTCTTTGCT-3’; R primer: 5’-CAGCATGGTAAGTGTGTTTCG-3’). PCR products were purified using the Qiaquick PCR Purification Kit (Qiagen). The EnGen Mutation Detection Kit (NEB) was used to identify mosaicism for *csf2rb* mutations. Once T7 digestion indicated the presence of CRISPR/Cas9-induced mutations, injected fish were added to the system and raised to adulthood. At 3 months of age, CRISPR F0 fish were outcrossed against wild-type AB zebrafish, and pooled resultant larvae were screened for mutations as previously described. Clutches positive for mutations in *csf2rb* were added to the system and raised to adulthood. At 2–3 months of age, F1 fish were fin-clipped. PCR and gel electrophoresis were used to identify zebrafish heterozygous for a deletion mutation in *csf2rb*. gDNA from mutant heterozygotes was amplified using PCR and sequenced by Sanger sequencing (Genewiz). Fish that were predicted to carry the same mutation became founders for the mutant line and were in-crossed. Individual resultant embryos were sequenced with Sanger sequencing (Genewiz) to confirm the mutant sequence. The resulting mutation is a single base pair insertion in exon 3 (NC_007114.7:g.24346850_24346851insG). This results in a frameshift mutation predicted to yield a substitution of 14 amino acids followed by an early stop codon (NP_001107205.1:p.Val86GlyfsX15). These mutants were assigned the allele name ‘mss15’ by The Zebrafish Information Network (ZFIN).

### Chemical treatment of zebrafish

Single DSS treatment of zebrafish larvae was carried out as per published protocols^15^. 7 days post-fertilization zebrafish larvae were treated with 0.25% (w/v) colitis grade DSS (36,000-50,000 MW, MP Biomedical) for 24 hours with or without 350nM recombinant human (rh)GM-CSF (R&D Systems). Recovery conditions consisted of the removal of DSS and returning of zebrafish to egg water with or without 350nM rhGM-CSF for 5 hours.

### Lck-GFP^+^ cell isolation and scRNA-seq

Lck-GFP+ cells were isolated from larval zebrafish intestines using the MACSQuant Tyto Cell Sorter (Miltenyi Biotec). Cells were suspended in 1 mL PBS + 0.5% BSA, sorted at 4 mL/hr (<150 Pa), and diluted to 50 µL. Cells were counted (Nexcelom Auto 2000), and 40 µL of sorted cells were loaded into the 10X Genomics Chromium controller with the Single-Cell A Chip Kit. cDNA libraries were prepared using the Chromium Single-Cell 3′ Library, Gel Bead Kit v2.

### Larval zebrafish gut length measurement and neutral red quantification

Larval zebrafish intestines were measured and neutral red staining was performed as per published protocols^15,16^. The ratio of intestine length to body length was used to account for individual growth rate variations. Larval zebrafish were imaged using Nikon SMZ745 scope. The zebrafish gut was selected using ImageJ and average intensity of neutral red staining was measured.

### Larval zebrafish scRNA-seq

scRNA-seq on larval zebrafish intestines was performed according to previously published protocols^16^. N = 30 larval zebrafish intestines were dissected per condition. 10,000 cells were loaded onto the 10X Genomics Chromium controller with the Single-Cell A Chip Kit (10X Genomics). The cDNA libraries were generated using the Chromium Single-Cell 3′ Library, Gel_Bead_Kit_v2 (10X Genomics). Libraries were sequenced on Illumina Novaseq. The raw (unfiltered) output matrices were used for the clustering and downstream analysis in R package Seurat v4.0 (X). Data were filtered to include cells with 150 < features (unique genes) < 3500, and mitochondrial transcripts < 30%. Samples were individually normalized, and the 2000 most variable genes were identified for each sample. ‘FindIntegrationAnchors’ was used to integrate all zebrafish samples. The data were scaled and dimensionality reduction was performed. For integrations of zebrafish samples, the first 15 principal components were used to generate clustering, with a resolution of 0.8. Enriched genes for each cluster were identified using the ‘FindAllMarkers’ function. The default Wilcoxon Rank Sum test was used to determine significance, with a cutoff of Log_2_(fold change) = 0.25.

### Ingenuity pathway analysis on DEGs

Marker genes DGE was calculated for ILC3 versus ILC1 as described in the methods above. Differentially expressed genes, log-fold changes, and p values were imported into ingenuity pathway analysis (IPA) for all data sets (https://digitalinsights.qiagen.com/IPA) Core analysis was conducted on each data set using log-fold change values.

### Differential gene expression (DGE) analysis

DGE between clusters and samples was determined using the ’FindMarkers’ function in Seurat v4.0^23^, with the Wilcoxon Rank Sum test for significance.

### Gene expression visualization

To visualize gene expression across ILC clusters and condition in a heatmap, the plot heatmap function was used from the R package Scillus (https://github.com/xmc811/Scillus).

### Single-cell velocity analysis

RNA velocity was analyzed across four conditions using Velocyto (v0.17) to generate loom files, which were merged with Seurat objects and processed in scVelo (v0.2.4) to model transcription dynamics^24^. Cell velocities indicated gene expression changes, and phase plots highlighted ILCs per treatment group.

### RNAscope in-situ hybridization

Fixation, cryosectioning, and immunofluorescence of larval zebrafish followed established protocols^25^. Larvae were fixed in 4% PFA overnight at 4°C, incubated in 30% sucrose, embedded in OCT, and stored at -80°C. Sections were mounted on VWR® Premium Superfrost® Plus slides. RNAscope ISH (ACD Bioscience) used four probe sets targeting csf2rb, il7r, rorc, ifng1.1, il22, tnfa, tbx21, il1b, and osm. Negative controls omitted probes. Sections were post-fixed, dehydrated, treated with Protease IV, and hybridized with diluted probes. Fluorophores identified C1-C4 targets. DAPI stained nuclei, and slides were mounted in EcoMount. Imaging (Zeiss 880, x20/x40) was standardized, with ≥5 zebrafish per condition. ImageJ was used for processing and quantification.

### Immunohistochemistry

Fixation, cryosectioning, and immunofluorescence of larval zebrafish were performed as described in the RNAscope in situ hybridization section. Cryosections (10 μm) were incubated overnight at 4°C with pSTAT5 (Tyr694) antibody (Invitrogen) at a dilution of 1:25 (2 μg), followed by incubation with Alexa Fluor 555-conjugated anti-mouse IgG (1:500; Invitrogen). Nuclei were counterstained with DAPI, and slides were mounted using EcoMount. Imaging was performed using a Zeiss 880 microscope at 40× magnification and standardized across samples, with at least 10 zebrafish per condition. ImageJ/Fiji was used for image processing and quantification.

Human tissue specimens were fixed in 10% formalin, paraffin-embedded, and sectioned at 3μm. Highplex IF was performed on the VENTANA-DISCOVERY-ULTRA (Roche Diagnostics) for automated processing. Adjacent sections were incubated with 1:6 CD14-PE (R&D) and DAPI or 1:6 GM-CSF-Alexa405, 1:30 pSTAT3-Alexa488, 1:30 pSTAT5-PE, and 1:30 pERK1/2-Alexa647 (Cell Signaling) for 1 hour at RT. Slides were mounted with Vectashield (Vector). Images were acquired at 300X using a Leica Stellaris 8 confocal microscope, merging 200-380 tiles per section. Pearson correlation coefficients were calculated using Coloc 2 (ImageJ/Fiji).

### Quantification and statistical analysis

Data points represent biological replicates, consisting of the mean of an individual patient or animal (as detailed in the figure legends) ± SEM. Statistical analyses were performed using GraphPad Prism (v9-11; GraphPad Software). For comparisons involving two groups, statistical significance was determined using the two-tailed Mann–Whitney U (Wilcoxon rank-sum) test. For experiments involving more than two groups, statistical significance was assessed using the Kruskal–Wallis test followed by Dunn’s multiple-comparisons test. Differential gene expression analyses of single-cell RNA-seq datasets were performed using the Wilcoxon rank-sum test implemented in Seurat v4.0. Ligand–receptor interaction analyses in stLearn used permutation testing with Benjamini–Hochberg false discovery rate (FDR) correction as described above. Statistical methods specific to each analysis are described in the corresponding Methods sections and figure legends.

### Materials availability

The *csf2rb^mss^*^15^ allele information is available at The Zebrafish Information Network (https://zfin.org/ZDB-ALT-220708-1#summary). Requests for embryos can be made to Ling-shiang Chuang and Jaime Chu.

## Results

### Differential myeloid cell growth factor abundance by inflammation and source-target mappings defined by spatial transcriptomics in ileal Crohn’s disease

We performed Xenium *In Situ* transcriptome profiling on paired inflamed and uninvolved ileal resection tissues from ten CD patients comparing M-CSF (CSF1), GM-CSF (CSF2) and G-CSF (CSF3). We observed markedly higher expression of M-CSF compared to GM-CSF and G-CSF (**Figure 1A-B**). underscoring its role in maintaining myeloid cell homeostasis in the gut^26,27^. Additionally, GM-CSF was uniquely enriched in inflamed vs. uninvolved regions across paired patient samples (**Figure 1C**), indicating its selective association with intestinal inflammation. Although GM-CSF-positive cells represented only a small fraction of total cells, even within inflamed tissue, this observation is consistent with the biology of GM-CSF as an inducible cytokine that is transiently produced by activated immune cells. In contrast to constitutively expressed growth factors such as M-CSF, the focal distribution of GM-CSF suggests that its spatial localization may be particularly relevant to its function within the inflamed tissue microenvironment.

**Figure 1.**
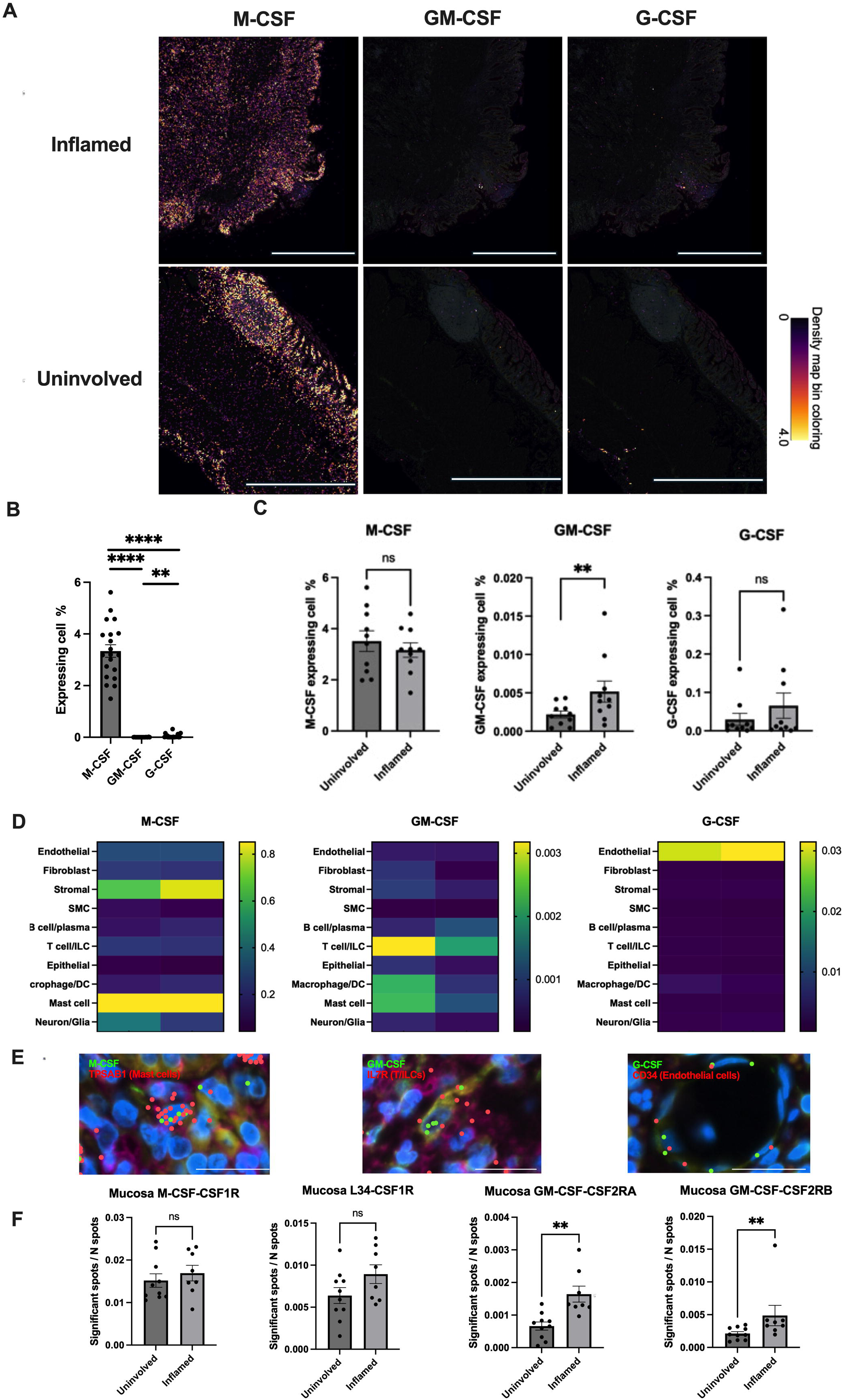
Elevated and localized GM-CSF expression in inflamed Crohn’s Disease ileums with stable M-CSF and G-CSF levels. **A.** Ileal Crohn’s disease tissue sections (inflamed and uninvolved regions) were profiled using 10X Xenium. Transcript density maps for M-CSF, GM-CSF, and G-CSF are color-coded: yellow (high), purple (intermediate), and black (none). Scale bar, 2000 µm. **B.** Bar graph quantifying the percentage of cells expressing each CSF per Xenium slide from 10 patients (10 inflamed and 10 uninvolved slides). **C.** Bar graph compares CSF expressions between inflamed and uninvolved regions (N = 10, **, P < 0.01). **D.** Heat map of CSF expression per cell cluster (yellow: high, green: medium, purple/black: low/none). **E.** Images of CSF with markers: TPSAB1 (mast cells), IL7R (T/ILC), and CD34 (endothelial cells). Scale bar, 20 μm. **F.** Bar graphs showing spatially inferred ligand-receptor interactions in inflamed vs. uninvolved ileal sections. The significant Spot to N spot sample ratio was calculated as the number of ligand-receptor interactions in the same grid (Spot) relative to the total spots in the full mucosa area (10 uninvolved slides and 8 inflamed slides). **, P < 0.01.

Xenium datasets revealed expression of M-CSF predominantly in stromal and mast cells and G-CSF in endothelial cells, with minimal differences due to inflammation status (**Figure 1D, Figure S1**). In contrast, GM-CSF demonstrated the greatest per cell expression in lymphocytes (T cells/ILCs) in inflamed tissues, with more modest per cell expression in myeloid cells (macrophages, dendritic cells, and mast cells). The cellular sources of CSFs identified by Xenium spatial transcriptomics were validated using published single-cell RNA sequencing data from Martin *et al.* Consistent with our findings, CSF2RB, the β-chain of the GM-CSF receptor, was broadly expressed across multiple cell types, including endothelial cells, B/plasma cells, monocytes/macrophages, dendritic cells (DCs), and mast cells. In contrast, the co-receptor CSF2RA displayed a restricted expression pattern, being detected exclusively in monocytes/macrophages and dendritic cells, indicating that these populations as candidate cellular targets of GM-CSF signaling within the intestinal microenvironment (**Figure S2A-B).** Representative high-magnification Xenium images further confirmed the focal enrichment of GM-CSF (CSF2) within inflamed tissue, while demonstrating the spatial distribution of CSF1, CSF3, and their corresponding receptors (CSF1R, CSF2RA, and CSF2RB) in uninvolved and inflamed ileal sections (**Figure S2C**). Spatial analysis using stLearn revealed that the ratio of significant GM-CSF ligand–receptor interaction spots (GM-CSF with either CSF2RA or CSF2RB) to total spots was significantly increased in inflamed compared to uninvolved intestinal tissue sections. This pattern was distinct from that observed for M-CSF and its receptor interactions, which showed no comparable change between conditions (**Figure 1F**). The enrichment of GM-CSF together with increased spatial proximity of GM-CSF and its receptor-expressing cells during inflammation, combined with genetic associations involving CSF2RB (including a frameshift, loss-of-function allele^5,28^) motivated further investigation of GM-CSF signaling during intestinal inflammation.

### Human GM-CSF mitigates intestinal inflammation in zebrafish in a Csf2rb-dependent manner

To test the functional role of the GM-CSF pathway during intestinal inflammation *in vivo*, we generated a novel zebrafish knockout line for *csf2rb*, the zebrafish ortholog of human *CSF2RB*. To generate *csf2rb*-deficient animals, we injected zebrafish embryos with CRISPR-Cas9 complex comprised of CRISPR RNA targeting *csf2rb* exon 3, fluorescently-labeled trans-activating CRISPR RNA, and Cas9 protein. Fluorescent-positive embryos, indicating successful delivery of CRISPR-Cas9 (**Figure S3A**), acquired mutations in the target region as assessed by T7 endonuclease I digest. Sequencing analysis identified a single-nucleotide (guanine) insertion in exon 3 of *csf2rb*, which was predicted to result in a frameshift that induces an early stop codon, yielding a truncated form of Csf2rb consisting of 99 amino acids (**Figure S3B-C**).

To distinguish the effects of GM-CSF during active inflammation from those during tissue repair, we employed two complementary experimental paradigms. In the injury model, zebrafish larvae were co-treated with DSS and recombinant human GM-CSF for 24 hours. In the recovery model, DSS was removed after 24 hours of injury, and larvae were subsequently allowed to recover for 5 hours in the presence or absence of GM-CSF (**Figure 2A**). Co-treatment with GM-CSF significantly reduced intestinal injury as measured by partial rescue of lysosome-rich enterocyte uptake of neutral red, a proxy for assessing epithelial cell function^29,30^, and rescued gut length, with loss of Csf2rb completely abrogating rescue by GM-CSF during DSS-induced injury (**Figure 2B-C**). In contrast, GM-CSF did not produce a detectable improvement during the short recovery period, suggesting that its protective effects in this model are most evident during active DSS-induced intestinal injury in vivo.

**Figure 2.**
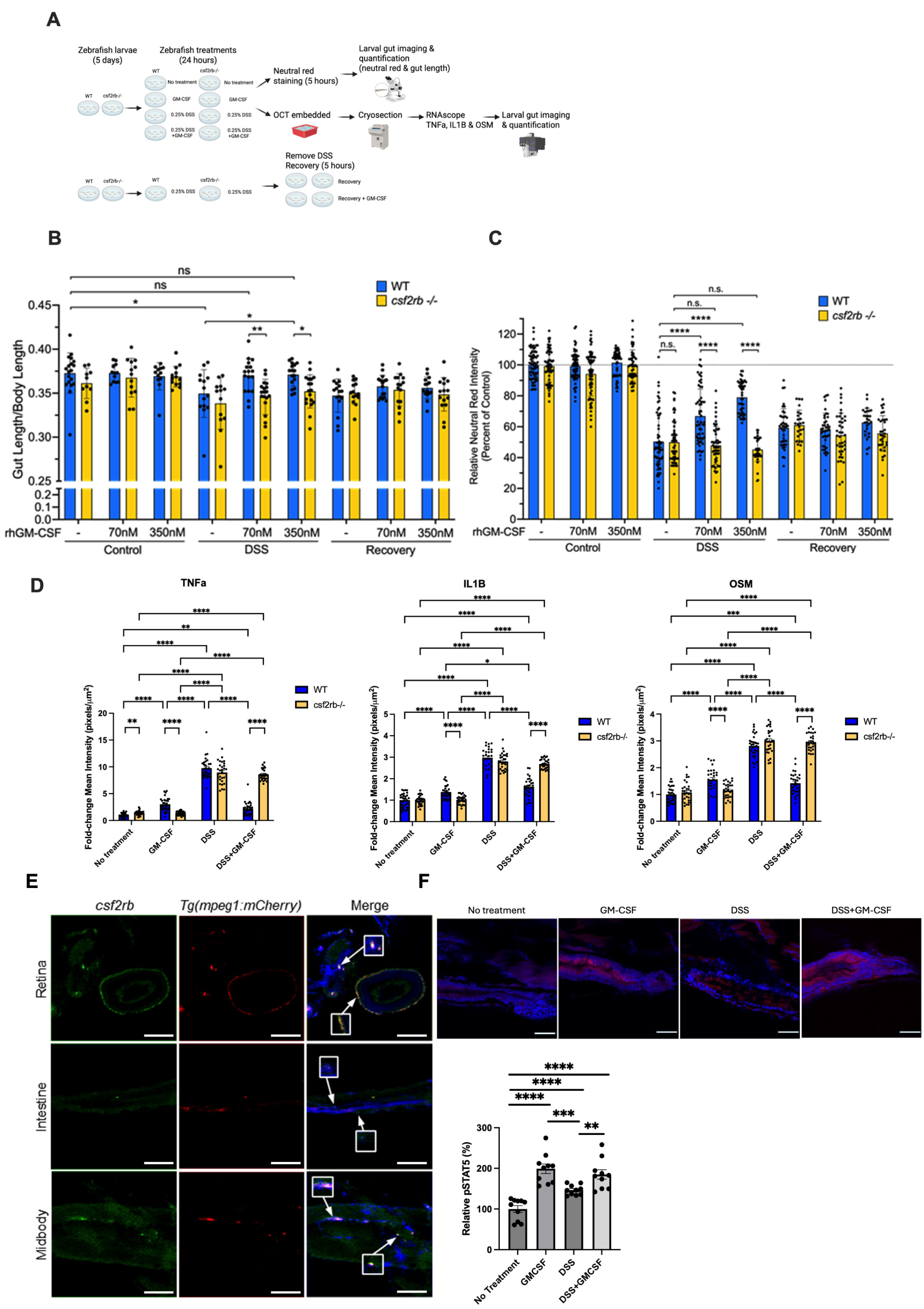
Human GM-CSF prevents injury in zebrafish DSS model of intestinal injury through Csf2rb signaling. **A.** Diagram of strategies on zebrafish treatments and data collections. **B.** Bar graph of intestinal length relative to total body length for WT (blue, N = 12-19/condition) and *csf2rb^-/-^* (yellow, N = 10-19/condition) larval zebrafish treated with DSS and GM-CSF. *, P < 0.05, **, P < 0.01. **C.** Bar graph of neutral red staining in the zebrafish intestine relative to genotype-matched controls for WT (blue, N = 28-70/condition) and *csf2rb^-/-^* (yellow, N = 25-68/condition) larval zebrafish treated with DSS and GM-CSF. ****, P < 0.0001. **D.** Quantification of RNA foci of RNAscope images for the proinflammatory cytokines *tnf*α, *il1b*, and *osm* in larval zebrafish intestines treated with DSS and human GM-CSF (*N* = 5, six images per larval gut). **, *P* < 0.01; ****, *P* < 0.0001. **E.** RNAscope detection of csf2rb transcripts (green) in *Tg(mpeg1:mCherry)* larval zebrafish, in which macrophages are labeled with mCherry (red). Representative csf2rb-positive mpeg1:mCherry-positive macrophages in the retina, intestine, and midbody are highlighted by arrows and magnified in the boxed insets. Additional csf2rb-positive cells lacking mpeg1:mCherry signal are also present, indicating that csf2rb expression is not restricted to macrophages. Scale bar, 100 µm. **F.** Immunostaining of phospho-STAT5 (red) and nuclei (blue, DAPI) with wildtype larval zebrafish treated with DSS and GM-CSF. Scale bar: 50 µm. **G.** Quantification of relative intensity of phospho-STAT5 per larval zebrafish gut. N = 10 per treatment conditions. **, *P* < 0.01; ***, P>0.001; ****, *P* < 0.0001.

To determine how GM-CSF further modulated the immune signaling environment of the inflamed intestine, we assessed expression of key cytokines in inflammation by RNAscope. RNA foci of pro-inflammatory cytokines *tnfa*, *il1b*, and *osm* were all induced with DSS treatment regardless of genotype; however, GM-CSF significantly blunted this increase specifically in WT zebrafish (**Figure 2D, S4A**). Importantly, in WT zebrafish, GM-CSF alone did induce pro-inflammatory gene expression, albeit at much lower levels than DSS treatment. However, these inductions of proinflammatory cytokines were abrogated in *csf2rb^-/-^* zebrafish (**Figure 2D, S4A**). To validate that *csf2rb* expression is conserved in zebrafish macrophages, we performed RNAscope in situ hybridization in *Tg(mpeg1:mCherry)* larvae, a transgenic line that labels macrophages with mCherry. Representative *csf2rb* RNA foci colocalized with *mpeg1:mCherry*-positive macrophages in the intestine, retina, and midbody (**Figure 2E, arrows and insets, S4B**), confirming that intestinal macrophages express *csf2rb*. Additional *csf2rb**-***positive cells that did not colocalize with mpeg1:mCherry were also observed, consistent with the broader expression pattern of CSF2RB identified in our human Xenium dataset. Thus, these data demonstrate that macrophages express *csf2rb*, but do not imply macrophage-restricted expression.

To assess STAT5 activation following recombinant human GM-CSF treatment in the zebrafish gut, pSTAT5 immunostaining was performed on WT zebrafish following GM-CSF and DSS treatment conditions. pSTAT5 levels were significantly increased with GM-CSF, DSS, and GM-CSF+DSS treatments compared to untreated controls, with the largest increase observed following exogenous GM-CSF administration (**Figure 2F-G**). These findings demonstrate increased STAT5 phosphorylation following GM-CSF exposure, consistent with activation of the GM-CSF signaling pathway in the zebrafish intestine.

### ILC-like populations with shared transcriptional features are identified in the larval zebrafish gut

Given that ILCs are potent expressors of GM-CSF in the inflamed human intestine and demonstrate dynamic population regulation, with reduced ILC3s and expanded ILC1s reported in inflamed CD patient tissues^4^, we reasoned that ILC-associated responses might contribute to the effects of GM-CSF during intestinal inflammation. Given that GM-CSF is largely produced by lymphocytes during intestinal inflammation, To this end, we characterized the lymphocyte populations present in the larval zebrafish intestine by dissecting intestines from Tg(lck:lck-GFP) zebrafish (N = ∼200/condition), a reporter line labeling lymphocytes with GFP, isolated GFP^+^ cells, and performing scRNA-seq (**Figure 3A**). Unsupervised clustering identified three major lymphocyte populations with ILC-associated transcriptional features (**Figure 3B**). Using this FACS-isolated intestinal lck:GFP+ lymphocyte scRNA-seq dataset, we examined the expression of previously reported zebrafish ILC markers (*nitr4a, nitr5,* and *nitr9*)^17^, conserved ILC3-associated genes (*il7r, rorc, il23r,* and *ccl20a.3*)^10^, and the proliferation marker *mki67* across the three clusters (**Figure 3C**). Based on these markers, together with conserved transcriptional regulators and effector programs, we annotated the three populations as proliferative ILC-like cells, putative ILC1-like cells, and putative ILC3-like cells (**Figure 3C-D**). The proliferative ILC population expressed proliferation markers such as *mki67* and *top2a* as well as the DNA methyltransferases *dnmt1*, *dnmt3bb.1* and DNA-associated components including *hmgn2*, *hmga1a*, and *h2az2b*. The putative ILC1-like population was characterized primarily by enrichment of a stress-response transcriptional module, including heat shock protein family members such as *hsp70l*, *hsp90aa1.2*, and *dnajb1b* (**Figure 3D**). Although similar stress-associated genes have been reported in human intestinal ILC1 datasets, these genes are not lineage-specific and therefore do not independently establish ILC1 identity. Accordingly, we interpret this population as a putative ILC1-like transcriptional state rather than a definitive zebrafish equivalent of mammalian ILC1s^17,31^. The putative ILC3-like population showed expression of conserved ILC3-associated transcriptional regulators, including *mafb*, *tcf7*, and *rorc*^32,33^ (**Figure 3D**).

**Figure 3.**
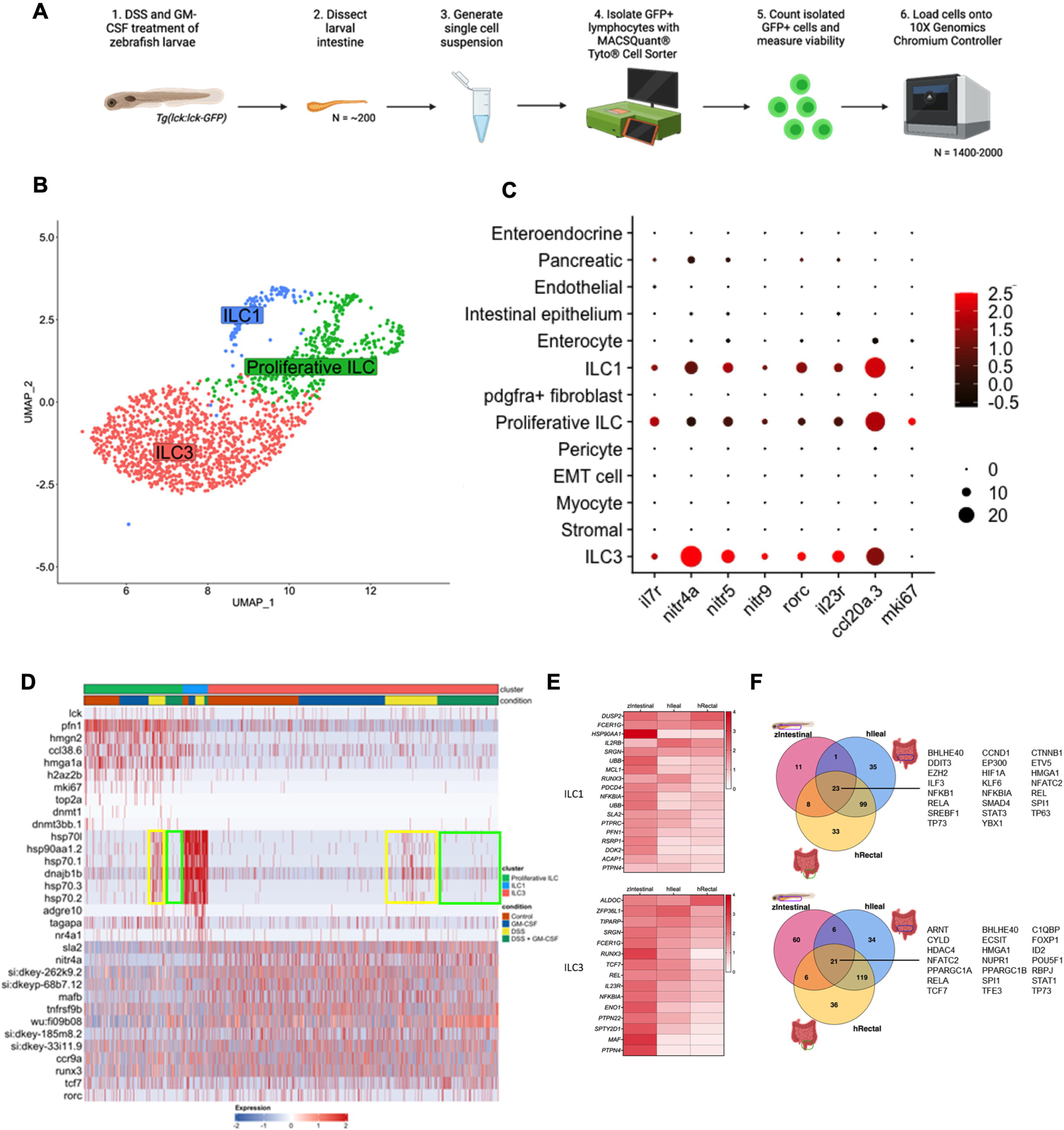
Single-cell transcriptomic profiling identifies zebrafish intestinal ILC-like populations with shared transcriptional features. **A.** Diagram of strategies on isolation and single-cell analysis of innate lymphoid cells (N = 200 larval guts/condition). **B.** UMAP visualization of intestinal lck:GFP-positive lymphocytes and transcriptionally annotated ILC-like populations from larval zebrafish. **C.** Dot plot generated from the scRNA-seq dataset of FACS-isolated intestinal lck:GFP+ lymphocytes from larval zebrafish, showing expression of conserved ILC3 markers (rorc and il23r), previously reported zebrafish ILC markers (il7r, nitr4a, nitr5, nitr9, and ccl20a.3), and the proliferation marker mki67 across the identified ILC-like clusters. **D.** Top differentially expressed genes for zebrafish proliferative ILC-like cells (top bar, green), ILC1-like cells (top bar, blue), and ILC3-like cells (top bar, orange) in no treatment control (bottom bar, red), GM-CSF (bottom bar, blue), DSS (bottom bar, yellow) and DSS GM-CSF cotreatment (bottom bar, green). The yellow rectangle highlights the increase expression of putative ILC1-like transcriptional module in proliferative ILC-like and putative ILC3-like populations following DSS treatment, whereas the green rectangle indicates attenuation of this transcriptional response following cotreatment with human GM-CSF. **E**. Heatmap showing shared transcriptional features between zebrafish putative ILC1-like (top panel) and ILC3-like (bottom panel) populations and human intestinal ILC subsets from human ileum and rectum tissue. **F.** Venn diagram showing overlapping predicted upstream transcriptional regulators identified by Ingenuity Pathway Analysis (IPA) for zebrafish putative ILC1-like and human ILC1 populations (top panel) and zebrafish putative ILC3-like and human ILC3 populations (bottom panel).

To explore the similarity of zebrafish and human ILCs, we compared the marker gene profiles of putative zebrafish ILC1-like and ILC3-like populations with human intestinal ILC subsets using zebrafish intestinal, human ileal^19^ and rectal^34^ scRNA-seq datasets. The zebrafish putative ILC1-like population and human ILC1s shared expression of HSP90AA1, UBB, MCL1, and DOK2, representing overlapping stress-response and cellular survival-associated transcriptional features rather than ILC1-specific lineage markers. The putative ILC3-like population shared expressed TCF7, IL23R, MAF, and PTPN22 with human ILC3s, consistent with partial conservation of ILC3-associated transcriptional programs. RUNX3, a transcription factor involved in ILC development and function, was enriched across both species (**Figure 3E**)^35^.

Ingenuity Pathway Analysis identified approximately 75% overlap among predicted upstream transcriptional regulators associated with the zebrafish putative ILC1-like and human ILC1 transcriptional profiles, including regulators associated with IL-2 and NF-κB signaling. The putative ILC3-like populations also showed overlap in predicted transcriptional regulators, including ARNT^36^, ID2^37^, RBPJ^38^, and TCF7^39^. Together, these analyses identify shared transcriptional features between zebrafish ILC-like populations and human intestinal ILC subsets, while acknowledging that transcriptional similarity alone does not establish equivalent lineage identity or function across species (**Figure 3F**).

### GM-CSF modulates ILC-like transcriptional states and inflammatory cytokine responses during intestinal inflammation

To assess changes in ILC-like populations during intestinal inflammation and GM-CSF signaling, we compared the relative proportions of the identified ILC-like populations across control, GM-CSF, DSS, and DSS + GM-CSF conditions in zebrafish intestines. DSS treatment resulted in a more than two-fold increase in the proportion of putative ILC1-like cells compared with controls, which returned toward control levels following GM-CSF co-treatment (**Figure 4A**). DSS treatment also resulted in a modest decrease in the proportions of proliferative ILC-like and putative ILC3-like cells, with the latter restored toward control levels following GM-CSF co-treatment (**Figure 4A**). To further evaluate these transcriptional changes, we used RNAscope *in situ* hybridization^40^ on larval zebrafish tissues following DSS +/-GM-CSF treatment. Expression of il7r, an ILC-associated marker^41^, remained relatively stable across conditions, suggesting that the overall abundance of il7r-expressing cells was not markedly altered (**Figure S4C-D**). To investigate whether changes in ILC-associated transcriptional markers were dependent on Csf2rb signaling, we quantified RNA foci corresponding to tbx21 and rorc, transcription factors associated with type 1 and ILC3-like programs, respectively. DSS treatment significantly increased tbx21 RNA foci regardless of genotype, whereas GM-CSF co-treatment attenuated this increase only in WT zebrafish (**Figure 4B-C, S4A**). Furthermore, DSS treatment decreased rorc RNA foci in WT zebrafish, an effect that was reversed by GM-CSF co-treatment (**Figure 4C**, right panel, **S4A)**. Together, these findings indicate that GM-CSF modulates inflammation-associated changes in tbx21 and rorc expression in a Csf2rb-dependent manner, consistent with the changes in ILC-like population proportions observed by scRNA-seq. Interestingly, proliferative ILC-like and putative ILC3-like populations exhibited increased expression of genes associated with the putative ILC1-like transcriptional state during intestinal inflammation (**Figure 3D**). To quantify this, we generated a putative ILC1-like transcriptional module from genes enriched in the corresponding zebrafish cluster and calculated the average expression across individual ILC-like cells. DSS treatment significantly increased the expression of this module in both proliferative ILC-like and putative ILC3-like populations, whereas GM-CSF co-treatment attenuated these changes (**Figure 4D**). These observations indicate inflammation-associated transcriptional changes but do not establish differentiation or lineage conversion between ILC-like populations.

**Figure 4.**
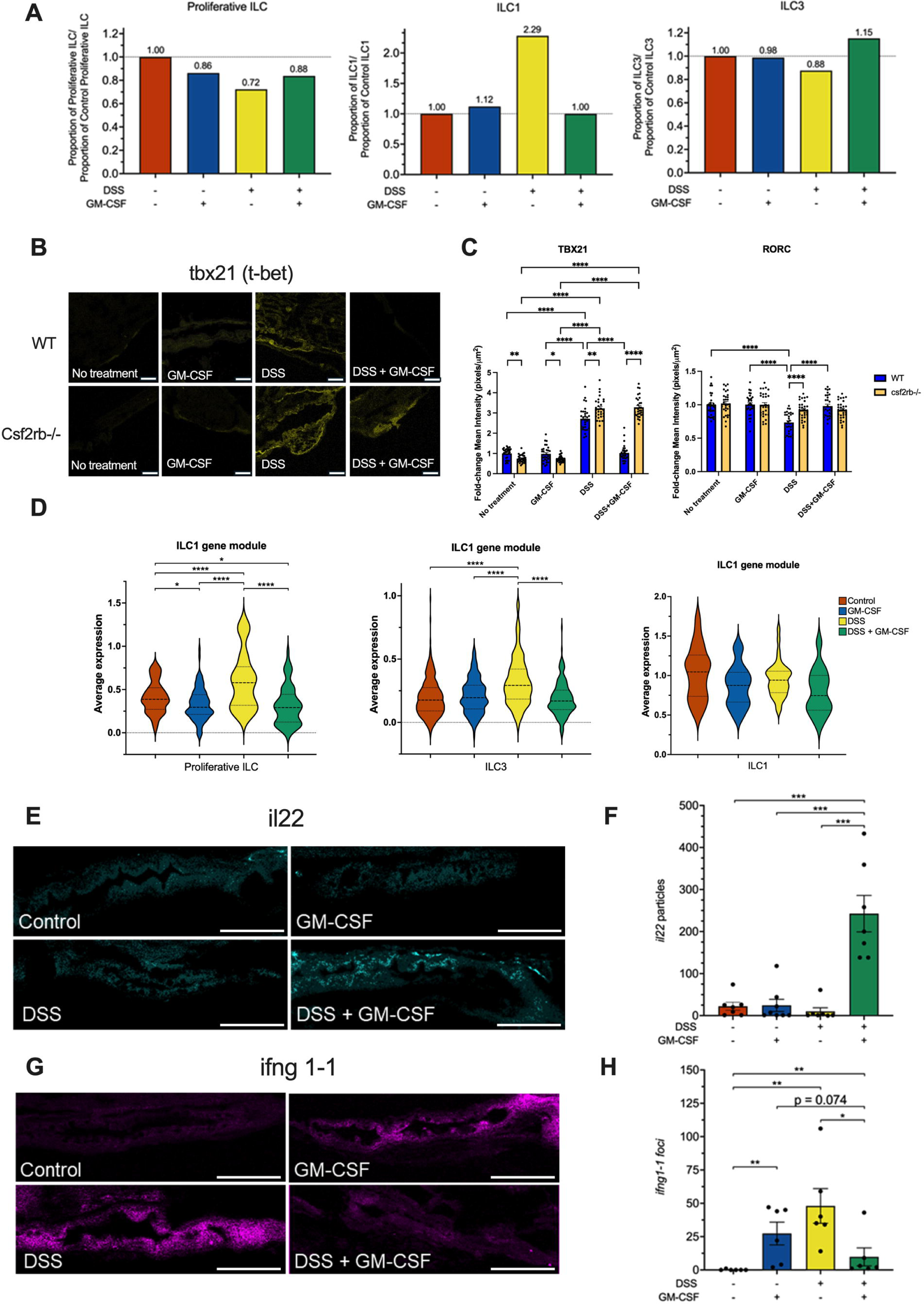
GM-CSF modulates ILC-like transcriptional states and inflammatory cytokine responses during intestinal inflammation. **A.** Relative proportions of transcriptionally annotated ILC-like populations in zebrafish intestinal scRNA-seq dataset across control, GM-CSF, DSS, and DSS + GM-CSF conditions. **B.** RNAscope images showing TBX21 probe staining in WT and csf2rb-/- zebrafish intestines treated with DSS and GM-CSF. Scale bar, 50 µm. **C.** Quantification of RNA foci for *tbx21* and *rorc*, transcription factors associated with type 1 and ILC3-like transcriptional programs, respectively (N = 5 zebrafish, six images per gut). *, P < 0.05; **, P < 0.01; ****, P < 0.0001. **D.** Expression of the putative ILC1-like transcriptional module in proliferative ILCs, putative ILC3-like cells, and putative ILC1-like cells under control, GM-CSF, DSS, and DSS + GM-CSF conditions (red: control, blue: GM-CSF, yellow: DSS, green: DSS + GM-CSF). Module scores were calculated using genes enriched in the putative zebrafish ILC1-like cluster. ****, P < 0.0001; *, P < 0.05. **E.** RNAscope images with il22 probe staining of zebrafish intestines treated with DSS and GM-CSF. Scale bar: 100 µm. **F.** Quantification of il22 RNA foci (N = 6-7, one image per gut). ***, P < 0.001. **G.** RNAscope images with ifng1-1 probe staining of zebrafish intestines treated with DSS and GM-CSF. Scale bar, 100 µm. **H.** Quantification of ifng1-1 RNA foci (N=6-7, one image per gut). **, P < 0.01; *, P < 0.05.

To further assess cytokine responses associated with intestinal inflammation, we evaluated expression of ILC-associated cytokines by RNAscope. *il22*, a cytokine associated with ILC3 function and intestinal epithelial barrier regulation^42^, was markedly induced following DSS and GM-CSF co-treatment, but not following either treatment alone (**Figure 4E-F, S4D**). Conversely, *ifng1.1*, a cytokine associated with type 1 inflammatory responses, was significantly induced by DSS treatment, and this increase was partially attenuated by GM-CSF co-treatment (**Figure 4G-H, S4D).** Although these cytokines are associated with distinct ILC effector programs, their expression cannot be attributed exclusively to specific ILC subsets based on tissue-level RNAscope analysis. Together, these findings demonstrate that GM-CSF treatment is associated with changes in ILC-like transcriptional states and cytokine expression during intestinal inflammation.

### GM-CSF modulates transcriptional dynamics of ILC-like populations during intestinal inflammation

Given that intestinal inflammation was associated with changes in ILC-like transcriptional states that were attenuated by GM-CSF treatment, we performed RNA velocity analysis on our zebrafish ILC-like scRNA-seq dataset to better characterize transcriptional dynamics across these populations. Enterocytes, representing a differentiated epithelial population, exhibited relatively low proportions of unspliced transcripts (**Figure 5A**). In contrast, proliferative ILC-like, putative ILC1-like, and putative ILC3-like populations exhibited higher proportions of unspliced transcripts relative to spliced transcripts (**Figure 5A**), consistent with differences in ongoing transcriptional activity between these populations^43^.

**Figure 5.**
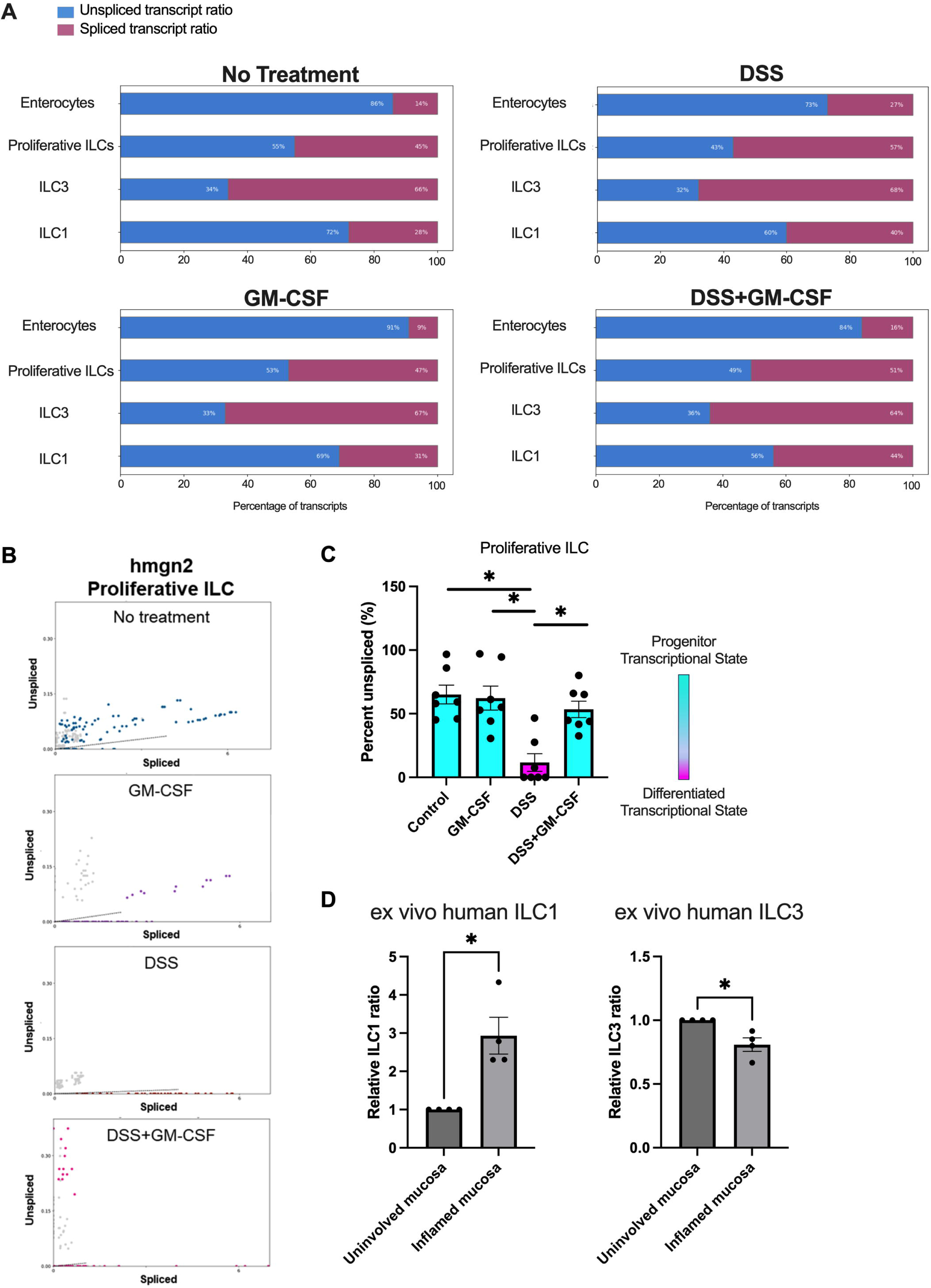
Transcriptional dynamics of intestinal ILC-like populations during inflammation. **A.** RNA velocity analysis showing the relative abundance of unspliced (red) and spliced (blue) transcripts in intestinal ILC-like populations and enterocytes in control (top left panel), GM-CSF (bottom left panel), DSS (top right panel) and DSS with GM-CSF co-treated (bottom right panel) larval zebrafish. **B.** Unspliced to spliced transcript ratios of proliferative ILC marker gene, *hmgn2* in control (top row, blue punctation), GM-CSF (second row, purple punctation), DSS (third row, red punctation) and DSS with GM-CSF co-treated (bottom row, pink punctation) larval zebrafish (N = 15/data point). **C.** Proportion of unspliced relative to spliced transcripts for the indicated proliferative ILC-like gene module. The y-axis provides the quantitative measure of unspliced transcript abundance, while the color scale visualizes the corresponding transcriptional state, with higher relative levels of unspliced transcripts indicating greater ongoing transcriptional activity rather than definitive differentiation potential. *, P < 0.05. **D.** Quantification of relative proportions of marker-defined ILC1-associated (IL7R+KLRB1+TBX21+) and ILC3-associated (IL7R+KLRB1+RORC+) populations in inflamed and uninvolved mucosa of human ileal tissue. *, P < 0.05.

The proliferative ILC-like population was characterized by expression of cell-cycle-associated genes, suggesting an actively proliferating transcriptional state^31^ rather than establishing a progenitor identity. One of its enriched markers, *hmgn2*, has been implicated in maintaining progenitor-associated chromatin states in other cellular contexts^44^. Analysis of the proportion of unspliced versus spliced *hmgn2* transcripts revealed that DSS-induced intestinal inflammation was associated with a reduced proportion of unspliced hmgn2 transcripts (**Figure 5B**). When we quantified these proportions for a module of top proliferative ILC marker genes, we similarly found a lower proportion of unspliced transcripts following DSS treatment. GM-CSF co-treatment restored these proportions toward control levels (**Figure 5C, S5A**). Together with the increased expression of the putative ILC1-like transcriptional module observed following DSS treatment, these findings suggest that intestinal inflammation alters transcriptional activity within proliferative ILC-like populations and that GM-CSF co-treatment attenuates these changes. However, these observations do not establish differentiation trajectories or lineage conversion between ILC-like populations.

Previous studies have demonstrated substantial plasticity among mammalian intestinal ILC populations, including the acquisition of ILC1-like characteristics by ILC3s under inflammatory conditions^43,45^. To examine whether inflammation was similarly associated with altered ILC subset representation in human intestinal tissue, we analyzed Xenium spatial transcriptomic data from paired inflamed and uninvolved ileal regions of CD patients. We observed an increased relative proportion of cells expressing the ILC1-associated marker combination (IL7R+ KLRB1+ TBX21+) and a decreased relative proportion of cells expressing the ILC3-associated marker combination (IL7R+ KLRB1+ RORC+) in inflamed compared with uninvolved tissue, consistent with previous reports^12,46^ (**Figure 5D, S5B**). Together, the zebrafish and human datasets demonstrate inflammation-associated differences in ILC-related transcriptional features and population composition. Although GM-CSF treatment attenuated transcriptional changes in zebrafish ILC-like populations, these findings do not establish direct regulation of ILC lineage identity or conversion.

### pSTAT5+ CD14+ macrophage aggregates exhibit spatial association with GM-CSF in the inflamed intestine

Intestinal ulceration is a common feature of CD inflammation^47^ and is histologically defined by disruption or loss of the epithelial barrier. The classification of tissue sections as inflamed or uninvolved was independently determined by pathological assessment of the resection specimens. Therefore, although uninvolved sections lacked the prominent inflammatory features characteristic of inflamed tissue, occasional focal epithelial breaks meeting the morphological criteria used for ulcer identification could still be observed (**Figure 6A-C**). In inflamed tissue, these epithelial disruptions were frequently associated with inflammatory cell-rich ulcer exudate and adjacent granulation tissue containing newly formed blood vessels^48^. While M-CSF exhibited widespread expression, GM-CSF and G-CSF were specifically expressed within ulcer granulation tissue in the inflamed ileum **(Figure 1A)**. Macrophage-rich aggregates were predominantly localized to the ulcer exudate, although macrophages were also occasionally observed within granulation tissue (**Figure 6B–F, S6A-B**). Interestingly, while inflammatory cytokines IL-1β and OSM were predominantly expressed within macrophage-rich exudate, TNF and GM-CSF were mainly localized to the granulation tissue (**Figure 6G–H, S6C**). These findings suggest that ulcer exudate and granulation tissue represent distinct inflammatory microenvironments with different spatial patterns of cytokine expression.

**Figure 6.**
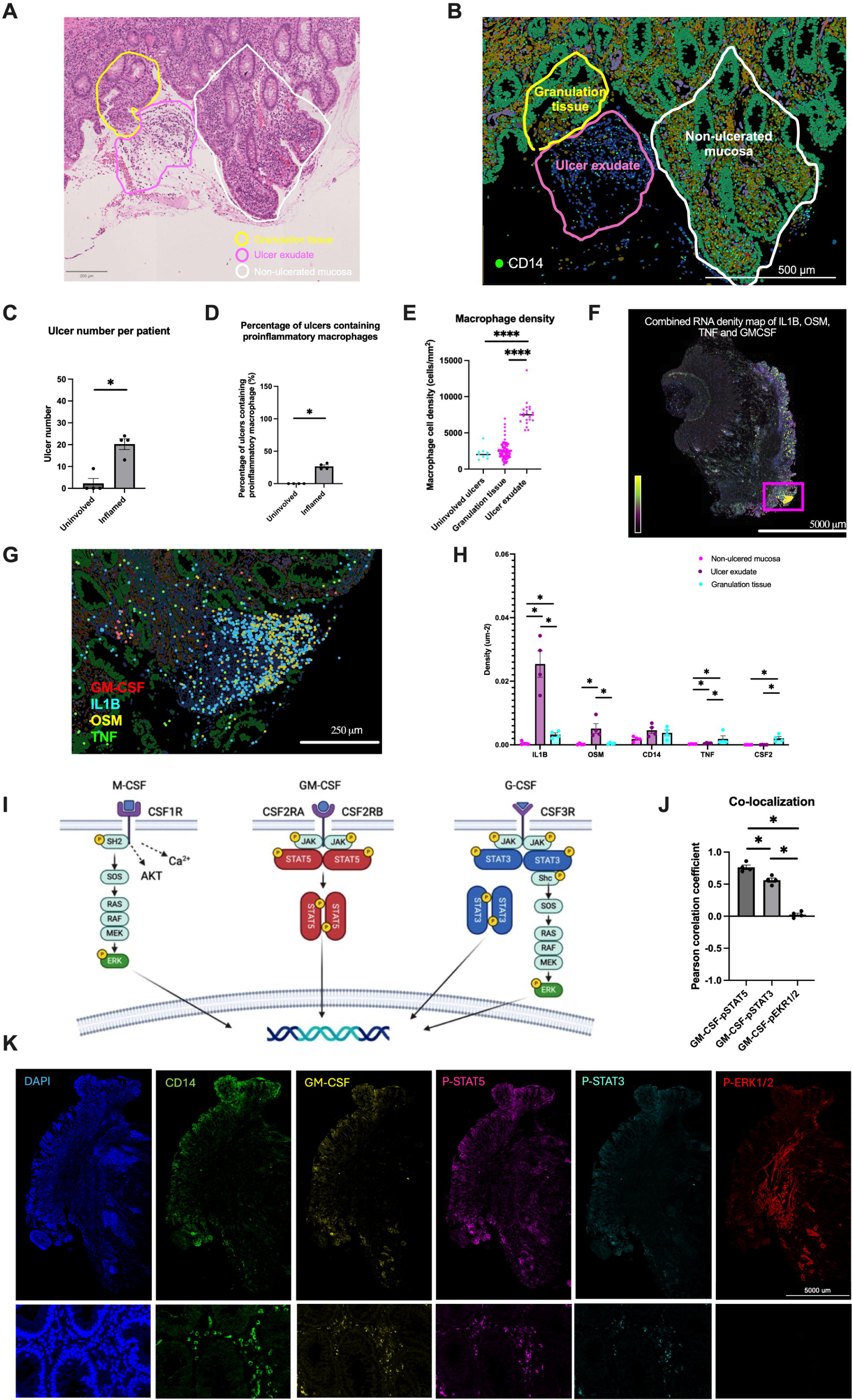
Spatial association of GM-CSF and pSTAT5 within macrophage-rich regions of inflamed Crohn’s disease intestine. **A.** H&E image of inflamed CD section with non-ulcerated mucosa (white), ulcer exudate (pink) and granulation (yellow). Scale bar, 200 μm. **B.** Xenium images of inflamed CD section with non-ulcerated mucosa (white), ulcer exudate (pink), and granulation tissue (yellow). Green cells represent the epithelial compartment and are distinct from the CD14-positive macrophage aggregates within the ulcer exudate. Scale bar, 500 μm. **C.** Quantification of ulcerative epithelial disruptions identified by loss or disruption of the epithelial barrier in pathologist-classified inflamed and uninvolved tissue sections. **D.** Quantification of the number of pro-inflammatory macrophage aggregates associated with ulcer exudate. **E.** Quantification of overall macrophage density within the defined tissue regions, independent of macrophage aggregate formation. * P < 0.05, **** P < 0.0001. N = 4 patients with paired inflamed and uninvolved sections. **F.** GM-CSF and cytokine (IL1B, TNF, OSM) density map. Scale bar, 5000 μm. **G.** Expression map of GM-CSF (red), IL1B (light blue), OSM (yellow), and TNF (green) showing IL1B and OSM enrichment in macrophage aggregates at ulcer exudate. Scale bar, 250 μm. **H**. Bar graphs of cytokine and CD14 transcript density in non-ulcered mucosa, ulcer exudate, and granulation tissue. *, P < 0.05. N = 4 patients with paired inflamed and uninvolved sections. **I.** Schematic illustrating canonical signaling pathways associated with M-CSF (ERK), GM-CSF (STAT5), and G-CSF (STAT3/ERK). **J.** Bar graph showing the Pearson correlation coefficient for GM-CSF and phosphorylated signaling proteins, calculated from five macrophage-rich regions per inflamed ileal section. **K.** Representative images of inflamed ileal tissue with co-immunostaining for DAPI, CD14, GM-CSF, p-STAT5, p-STAT3 and p-ERK1/2 (top panel). Scale bar, 5000 μm. Magnified views of macrophage-rich regions show DAPI, CD14, GM-CSF, p-STAT5, p-STAT3 and p-ERK1/2 (bottom panel). Scale bar, 250 μm.

Given the spatially restricted nature of GM-CSF, we aimed to explore if its downstream signaling occurred in a similar manner. STAT5 phosphorylation (pSTAT5) occurs downstream of GM-CSF engagement and multimerization of its receptors CSF2RA and CSF2RB. Unlike GM-CSF, M-CSF and G-CSF signal through ERK and STAT3 phosphorylation, respectively (**Figure 6I**). To assess CSF signaling spatially, we performed immunofluorescent staining for GM-CSF, pSTAT5, pSTAT3, and pERK1/2 on inflamed ileal sections, with DAPI and CD14 staining on adjacent sections. By quantifying the Pearson Correlation Coefficient across the analyzed macrophage-rich regions between GM-CSF and the phosphorylated signaling proteins, we observed a strong and significant colocalization between GM-CSF and pSTAT5 compared to pSTAT3 and pERK1/2 (**Figure 6J**). GM-CSF and pSTAT5 were localized to regions containing CD14-positive macrophage aggregates, consistent with previous transcriptomic findings (**Figure 6K and S6D**). In conclusion, local GM-CSF expression is associated with STAT5 phosphorylation within macrophage-rich regions, potentially contributing to tissue homeostasis during intestinal inflammation.

To further characterize the macrophage programs associated with GM-CSF signaling, we performed complementary transcriptomic analyses of human macrophages from a previously published Crohn’s disease scRNA-seq dataset and zebrafish macrophages under experimental GM-CSF treatment conditions. In the human dataset, STAT5-high macrophages, compared with STAT5-low macrophages, exhibited enrichment of STAT5A, STAT5B, and genes associated with immune regulation and tissue remodeling, including LGALS1, TGFBI, and GPNMB. Because these human macrophages were not experimentally treated with GM-CSF, these findings represent STAT5-associated transcriptional differences rather than a direct GM-CSF treatment response. In zebrafish, comparison of macrophages following DSS + GM-CSF versus DSS treatment identified differential expression of genes associated with retinoic acid signaling (rarab) and HIF pathway regulation (hif1a). Additional comparison of WT and csf2rb-deficient macrophages under GM-CSF treatment conditions identified Csf2rb-associated differences in these transcriptional programs (**Figure S7**). These findings identify candidate macrophage pathways associated with GM-CSF/Csf2rb signaling that may influence the intestinal immune environment during inflammation.

## Discussion

GM-CSF, M-CSF, and G-CSF direct myeloid cell differentiation, and myeloid cells are key drivers of inflammation and tissue damage in IBD^49^. Thus, clarifying the distinct roles of CSFs in CD pathogenesis remains essential. Genome-wide association studies (GWAS) have significantly advanced our understanding of inflammatory bowel disease (IBD). Our previous research identified a single nucleotide polymorphism (SNP) in *CSF2RB* as the second most prevalent frameshift SNP associated with CD in the Ashkenazi Jewish population, following *NOD2*^5^. Additionally, a recent study by Liu et al. in *Nature Genetics*^50^ reported a strong association between *CSF2RB* and CD in the East Asian population, with an exceptional P-value of 10^-20^, ranking third after *IL23R* and *ADO*. These findings highlight the genetic relevance of CSF2RB to CD susceptibility across distinct populations. Given that impaired GM-CSF signaling has been associated with increased CD susceptibility and complications^5,28^, GM-CSF likely plays a protective role in intestinal inflammation. In this report, we identify GM-CSF signaling niches comprised of innate lymphoid and myeloid cells that are induced specifically in granulation tissue of ulcers within inflamed ileal tissue from CD patients. We further show that recombinant human GM-CSF mitigates DSS-induced intestinal injury and attenuates pro-inflammatory cytokine expression (*il1b*, *osm*, *tnfa* and *ifng1-1)*, in a Csf2rb-dependent manner in zebrafish.

GM-CSF treatment was also associated with reduced proportions of putative ILC1-like cells, preservation of putative ILC3-like populations, increased il22 expression, and attenuation of inflammation-associated transcriptional changes in proliferative ILC-like cells (**Figure 7**).

**Figure 7.**
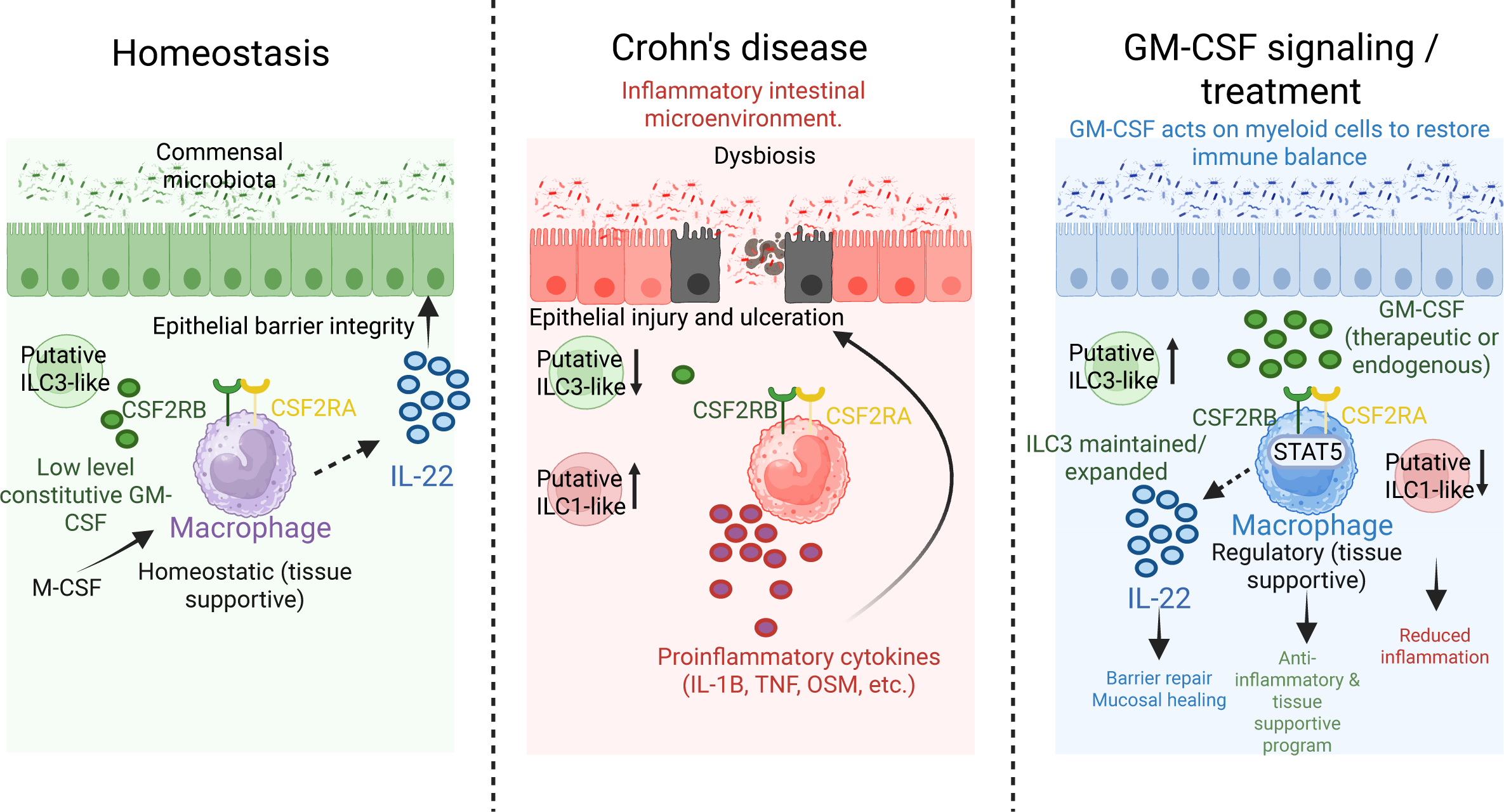
Proposed model of GM-CSF-associated myeloid and ILC-like responses during intestinal inflammation. Schematic illustrating M-CSF and GM-CSF signaling during intestinal homeostasis, Crohn’s disease inflammation, and GM-CSF treatment. GM-CSF acts primarily on myeloid cells through CSF2RA/CSF2RB-associated signaling. During intestinal inflammation, GM-CSF treatment attenuates epithelial injury and inflammatory cytokine expression, accompanied by preservation of putative ILC3-like populations, reduced putative ILC1-like representation, and increased IL-22 expression.

We leveraged Xenium *In Situ* single-cell spatial transcriptomics to characterize the distribution of CSF cytokines and their receptors in paired inflamed and uninvolved ileal tissues from CD patients. This fine-scale spatial resolution allowed us to identify that while M-CSF and G-CSF transcripts are abundant in the intestinal mucosa, GM-CSF was selectively enriched in inflamed tissue and exhibited focal localization within ulcer-associated regions. In contrast, M-CSF showed a relatively widespread distribution, and G-CSF is mainly confined to blood vessels. Notably, GM-CSF-expressing cells were rare even in inflamed tissue, consistent with prior reports that GM-CSF is tightly regulated and produced transiently by activated lymphoid and myeloid cells rather than constitutively across the mucosa. Unlike more broadly distributed growth factors such as M-CSF, the focal expression of GM-CSF is consistent with a model of localized paracrine signaling, in which GM-CSF-producing cells may influence neighboring CSF2RA/CSF2RB-expressing myeloid populations. The spatial association between GM-CSF and pSTAT5 within ulcer-associated macrophage-rich regions further supports the possibility of localized GM-CSF signaling during intestinal inflammation. These distinct spatial patterns suggest that GM-CSF may contribute to the organization of local inflammatory microenvironments in CD.

Interestingly, GM-CSF transcripts were predominantly localized within ulcer-associated granulation tissue, whereas the pro-inflammatory cytokines IL1B and OSM were enriched in macrophage-rich ulcer exudate. Because granulation tissue is associated with tissue repair following inflammatory injury, this spatial distribution raises the possibility that GM-CSF participates in local tissue-repair responses rather than exclusively promoting inflammation. Although GM-CSF-expressing cells were relatively rare, their focal distribution may still be biologically relevant within these specialized tissue microenvironments. Previous studies have identified intestinal ILC3s as important sources of GM-CSF^51^, while our spatial transcriptomic analyses identified lymphocytes as prominent GM-CSF-expressing populations in inflamed tissue. Together, these observations suggest that interactions between GM-CSF-producing lymphocytes and neighboring myeloid cells may contribute to the local immune environment of intestinal ulcers.

To characterize GM-CSF signaling during intestinal inflammation *in vivo*, we generated the first zebrafish *csf2rb* knockout line (*csf2rb^mss15^*). While *CSF2RB* expression is conserved in myeloid cells between humans and zebrafish, orthologs for *CSF2RA* and *CSF2* remain unidentified— even with support from the Zebrafish Information Network (ZFIN)—likely due to low cross-species gene sequence conservation. Many zebrafish cytokines share conserved protein structures and functional similarities with their mammalian counterparts despite low sequence homology. Our data show that recombinant human GM-CSF rescues DSS-induced intestinal injury in zebrafish, an effect that is abolished in *csf2rb* knockout animals, supporting a conserved Csf2rb-dependent response to GM-CSF. Consistent with the established biology of GM-CSF, csf2rb-deficient zebrafish displayed no detectable baseline intestinal abnormalities under the conditions examined, suggesting that Csf2rb signaling is not essential for maintaining the intestinal phenotypes assessed during homeostasis. In contrast, during DSS-induced intestinal injury, recombinant human GM-CSF protected against epithelial damage and gut shortening in wild-type but not *csf2rb*-deficient larvae, supporting a role for GM-CSF as a protective regulator during active intestinal injury. Consistent with activation of downstream signaling, recombinant human GM-CSF treatment also significantly increased intestinal pSTAT5 levels in zebrafish.

To further define the macrophage response to GM-CSF, we compared transcriptional programs across human Crohn’s disease and zebrafish macrophages. Human STAT5-high macrophages exhibited regulatory and tissue-remodeling signatures characterized by LGALS1, TGFBI, and GPNMB, whereas zebrafish GM-CSF-responsive macrophages demonstrated activation of conserved retinoic acid- and HIF-associated pathways, including *rarab* and *hif1a*. Loss of csf2rb attenuated these programs, supporting a conserved role for GM-CSF in establishing macrophage states associated with tissue homeostasis rather than persistent inflammation.

ILCs are dynamically regulated by inflammation with ILC1s expanding and ILC3s contracting in IBD^4,46^. Furthermore, the level of serum anti-GM-CSF autoantibodies correlates with increased ileal ILC1s in CD patients^4^, highlighting a potential role for GM-CSF in shaping intestinal ILC-associated responses. Intestinal ILCs have been previously profiled in human intestinal tissue, and ILC-like populations have also been identified in the adult zebrafish intestine^17,52,53^. To study the potential for GM-CSF to modulate ILC-associated responses, we identified and transcriptionally characterized zebrafish intestinal ILC-like populations based on the currently available zebrafish markers and conserved mammalian transcriptional programs. We acknowledge that the absence of fully validated lineage markers in zebrafish precludes definitive assignment of mammalian-equivalent ILC subsets, and have therefore interpreted these populations as putative ILC-like transcriptional states rather than direct equivalents of mammalian ILC subsets. Our analyses identified shared transcriptional features and regulators between larval zebrafish ILC-like populations and human intestinal ILC subsets, although these similarities do not establish equivalent lineage identity or function. The putative ILC3-like population was the most abundant ILC-like cluster in the larval zebrafish intestine. Consistent with this, the putative ILC3-like cluster was approximately 11.2 times more abundant than the putative ILC1-like cluster in our zebrafish scRNA-seq dataset, with il7r expression remaining relatively stable across treatment conditions (Figure 3F; Supplementary Figure 4C-D). Thus, the observed twofold increase in the relative proportion of putative ILC1-like cells during DSS injury occurred against a substantially larger baseline population of putative ILC3-like cells, which may help explain the comparatively modest proportional decrease in the latter.

Our zebrafish experiments further demonstrated that recombinant human GM-CSF modulates the relative abundance of ILC-like populations and their associated transcriptional features during intestinal inflammation. While this model lacks a mature adaptive immune system at the larval stage ^54^ and does not fully recapitulate the immune environment in CD, it provides the unique opportunity to examine innate immune responses without the contribution of mature adaptive lymphocyte populations. Furthermore, this avoids the need for ablation of immune compartments by *Rag1/2^-/-^*and antibody-depletion based methods, commonly employed in mice, that disrupt the native immune system and can also alter disease-relevant responses. However, to further strengthen the physiological relevance of our findings, future studies should examine human intestinal ILCs isolated from patient-derived tissue or intestinal explants. This would enable direct assessment of GM-CSF-mediated signaling in a human immune context and enhance the translational value of our zebrafish model.

Additionally, we identify a population of proliferative ILC-like population characterized by enrichment of cell-cycle-associated genes. In humans, ILC progenitors are abundant in the fetal intestine and decreased in the adult intestine^52,53^. Although the transcriptional features of this zebrafish population suggest active proliferation, they do not establish a progenitor identity. Our scVelo analysis identified a reduced proportion of unspliced transcripts of the proliferative ILC-like marker hmgn2 following DSS-induced intestinal injury, consistent with altered transcriptional dynamics during inflammation. *HMGN2* is known to play a role in proliferation, and is enriched in many progenitor cells^44^, although its expression alone is insufficient to define progenitor status. GM-CSF co-treatment attenuated the DSS-associated changes in unspliced transcript abundance and the expression of the putative ILC1-like transcriptional module, suggesting modulation of inflammation-associated transcriptional states rather than demonstrating lineage conversion. Given the predominant expression of GM-CSF receptor components in myeloid populations, these findings raise the possibility that GM-CSF influences ILC-like states indirectly through macrophage-associated signaling. Retinoic acid signaling^4,5^, suggested by the macrophage transcriptional analyses, represents one candidate pathway, although its functional involvement in ILC-like regulation remains to be established.

GM-CSF signaling is enriched with intestinal inflammation and exhibits a distinct spatial distribution within ulcerated tissue. Our spatial transcriptomic analysis reveals that GM-CSF, and proinflammatory cytokines TNF, IL1B, and OSM, are expressed in distinct niches within ulcers in CD. Previous studies have shown that the *CSF2RB* frameshift mutation associated with increased CD risk results in decreased pSTAT5 in response to GM-CSF in macrophages^5^. Furthermore, genetic variants involving *STAT5A* and *STAT5B* have been associated with CD^2,55^ and mice deficient in STAT5 tetramerization are hypersusceptible to DSS colitis^56^. We found that pSTAT5, but not pERK, exhibited strong spatial colocalization with GM-CSF within macrophage-rich aggregates, consistent with localized GM-CSF-associated signaling. Although *CSF3* and pSTAT3 also appear in the aggregates, pSTAT3 demonstrated significantly less colocalization with GM-CSF. Together, these findings highlight a spatial association between GM-CSF and pSTAT5 within macrophage-rich regions of inflamed Crohn’s disease tissue, distinct from the spatial patterns observed for pSTAT3 and pERK. This observation supports a model in which localized GM-CSF signaling may contribute to the inflammatory and tissue-repair responses occurring within intestinal ulcers. While GM-CSF transcripts were localized within ulcer granulation tissue, GM-CSF protein was detected in both granulation tissue and ulcer exudate. This difference may reflect the secretion and diffusion of GM-CSF from its producing cells, such that protein localization does not necessarily correspond directly to transcript distribution.

Importantly, our study integrates spatial transcriptomic characterization of human Crohn’s disease tissue with functional investigation of GM-CSF/Csf2rb signaling in a zebrafish model of intestinal injury. Numerous studies have highlighted proinflammatory roles for GM-CSF in intestinal inflammation^7,51,57^; however, our findings demonstrate that recombinant human GM-CSF can also exert protective effects during DSS-induced intestinal injury in a Csf2rb-dependent manner, highlighting the context-dependent functions of this cytokine. By maintaining the equilibrium between immune cell populations, GM-CSF ensures that inflammatory responses are kept in check, preventing excessive tissue damage and dysregulated immune responses that characterize CD. These protective effects were accompanied by changes in ILC-like population proportions and inflammatory cytokine expression, suggesting that GM-CSF may contribute to maintaining intestinal immune balance during inflammation. These findings highlight the potential importance of GM-CSF signaling in intestinal homeostasis and motivate further investigation of its therapeutic relevance to Crohn’s disease. By integrating human spatial transcriptomics with functional zebrafish modeling, our study provides new insights into the spatial organization of GM-CSF-associated signaling and its relationship with myeloid and ILC-like responses during intestinal inflammation.

## Ethics Statement for Human Participant and Animals

Patients were identified by screening surgical programs at Mount Sinai Hospital. Protocols were reviewed and approved by the Institutional Review Board (IRB) at the Icahn School of Medicine at Mount Sinai (HSM#14-00210). Detailed information regarding the participants and resection samples can be found in **Supplemental table 1**-**2**.

All zebrafish experimental procedures were approved (protocol #: IACUC-2015-0050 & IPROTO202100000045) by Mount Sinai Institutional Animal Care and Use Committee (IACUC) in accordance with NIH guidelines for the humane treatment of animals. Adult fish were maintained on a 14:10 light/dark cycle at 28°C. Fertilized embryos collected following natural spawning were cultured at 28.5°C in egg water (0.6 g/l Instant Ocean Sea Salt; Blacksburg) containing methylene blue (0.002 g/l). AB WT, *csf2rb^mss^*^15^, *Tg(lck:lck-GFP)^cz2Tg^* ^58^, *Tg(mpeg1:mCherry)^gl^*^23^ ^59^ zebrafish strain were used.

## Data and code availability

Single-cell RNA-seq data have been deposited at GEO and are publicly available as of the date of publication. All original code has been deposited and is publicly available as of the date of publication.

## Acknowledgements

The authors thank the patients who participated in this study and staff that assisted in their recruitment. We would like to thank Kenneth Frazer from ZFIN for his assistance with querying the zebrafish genome for putative Csf2rb co-receptor and ligand orthologs. Zebrafish were maintained and cultivated at the Zebrafish Shared Research Facility (Z-SRF) at the Icahn School of Medicine at Mount Sinai with assistance from Victoria Sclar. The Flow Cytometry CORE at the Icahn School of Medicine at Mount Sinai provided training, consultation, and technical assistant. Microscopy was performed at the Microscopy CORE at the Icahn School of Medicine at Mount Sinai. Supported by the National Institute of Health (R01 DK123758), Inflammatory Bowel Disease Genetics Consortium (U01 DK062422), The Helmsley Charitable Trust, The Sanford J. Grossman Charitable, Crohn’s and Colitis Foundation Visiting IBD Research Fellowship and Department of Genetics and Genomic Sciences Pilot Award at Icahn School of Medicine.

