## Supplementary material for "GM-CSF modulates conserved ILC-like transcriptional states and myeloid cell signaling during ulceration in Crohn’s disease": S figure 1-6 table 1: Supplement elife V2.pdf

**Supplementary figure 1.** The tables display the percentage of cells expressing CSF in each cell cluster. Four pairs of inflamed and uninvolved resection tissues were analyzed.

### M-CSF (CSF1)

| | Inflamed | | | | Uninvolved | | | |
|---|---|---|---|---|---|---|---|---|
| Endothelial | 0.3069 | 0.2182 | 0.3207 | 0.2961 | 0.2455 | 0.3171 | 0.323 | 0.2605 |
| Fibroblast | 0.1682 | 0.3048 | 0.3208 | 0.1218 | 0.2826 | 0.2687 | 0.2229 | 0.0607 |
| Stromal | 0.4932 | 0.5359 | 0.9066 | 0.6729 | 0.9248 | 0.8492 | 0.9656 | 0.5238 |
| SMC | 0.0778 | 0.0936 | 0.145 | 0.0577 | 0.0584 | 0.0903 | 0.0718 | 0.054 |
| B cell/plasma | 0.0641 | 0.1319 | 0.12 | 0.1408 | 0.157 | 0.1592 | 0.1435 | 0.2037 |
| T cell/ILC | 0.1799 | 0.2928 | 0.181 | 0.2437 | 0.1689 | 0.2227 | 0.1886 | 0.2387 |
| Epithelial | 0.0562 | 0.0375 | 0.0564 | 0.0751 | 0.0314 | 0.0404 | 0.0516 | 0.0439 |
| Macrophage/DC | 0.0963 | 0.1675 | 0.0998 | 0.1648 | 0.2131 | 0.1832 | 0.1309 | 0.196 |
| Mast cell | 1.4458 | 0.4667 | 1.1129 | 0.3816 | 1.7497 | 0.5173 | 0.7521 | 0.3834 |
| Neuron/Glia | 0.342 | 0.3701 | 0.5391 | 0.4035 | 0.2283 | 0.3449 | 0.2589 | 0.0746 |

### G-CSF (CSF3)

| | Inflamed | | | | Uninvolved | | | |
|---|---|---|---|---|---|---|---|---|
| Endothelial | 0.0166 | 0.0076 | 0.09 | 0.004 | 0.0094 | 0.0058 | 0.1097 | 0.001 |
| Fibroblast | 0.0002 | 0.0005 | 0.0013 | 0.0001 | 0.0004 | 0.0002 | 0.0006 | 0.0002 |
| Stromal | 0.0013 | 0.0005 | 0.0023 | 0.0003 | 0.0013 | 0.0004 | 0.0031 | 0.0002 |
| SMC | 0.0002 | 0.0003 | 0.0011 | 0.0001 | 0.0002 | 0.0002 | 0.0003 | 0.0001 |
| B cell/plasma | 0.0004 | 0.0005 | 0.0008 | 0.0007 | 0.0006 | 0.0005 | 0.0008 | 0 |
| T cell/ILC | 0.0007 | 0.0008 | 0.0014 | 0.0007 | 0.0004 | 0.0007 | 0.0014 | 0.0002 |
| Epithelial | 0.0003 | 0.0006 | 0.0007 | 0.0009 | 0.0006 | 0.0004 | 0.0004 | 0.0002 |
| Macrophage/DC | 0.0003 | 0.0019 | 0.0037 | 0.0051 | 0.0011 | 0.0018 | 0.0028 | 0.0003 |
| Mast cell | 0.0002 | 0.0008 | 0.0019 | 0.0005 | 0 | 0.0008 | 0.0021 | 0.0002 |
| Neuron/Glia | 0.0004 | 0.0005 | 0.0028 | 0.0006 | 0.0015 | 0.0005 | 0.0039 | 0.0001 |

### GM-CSF (CSF2)

| | Inflamed | | | | Uninvolved | | | |
|---|---|---|---|---|---|---|---|---|
| Endothelial | 0.0005 | 0.0006 | 0.0006 | 0.0004 | 0.0005 | 0.0004 | 0.0008 | 0.0002 |
| Fibroblast | 0.0004 | 0.0007 | 0.002 | 0.0001 | 0.0004 | 0.0002 | 0.0004 | 0 |
| Stromal | 0.0005 | 0.0006 | 0.0019 | 0.0007 | 0 | 0.0007 | 0.001 | 0.0005 |
| SMC | 0.0003 | 0.0002 | 0.0002 | 0.0001 | 0.0002 | 0.0003 | 0.0001 | 0.0001 |
| B cell/plasma | 0.0003 | 0.0007 | 0.001 | 0.0006 | 0.0013 | 0.0016 | 0.001 | 0.0006 |
| T cell/ILC | 0.0027 | 0.0044 | 0.0037 | 0.0019 | 0.0022 | 0.0014 | 0.0026 | 0.0018 |
| Epithelial | 0.0006 | 0.0007 | 0.0006 | 0.0012 | 0.0003 | 0.0005 | 0.0005 | 0.0003 |
| Macrophage/DC | 0.0025 | 0.0013 | 0.0013 | 0.0038 | 0.0004 | 0.001 | 0.0011 | 0.0006 |
| Mast cell | 0.001 | 0.0007 | 0.0066 | 0.0009 | 0 | 0.0014 | 0.0025 | 0.0006 |
| Neuron/Glia | 0.0005 | 0.001 | 0.0006 | 0.0004 | 0.0006 | 0.0005 | 0.0003 | 0.0001 |

**Supplementary figure 2.** Validation of CSF source and target cells using published single-cell RNA sequencing data from *Martin et al.* **A.** Source cells expressing M-CSF (left), GM-CSF (middle), and G-CSF (right) in *ex vivo* inflamed versus uninvolved Crohn’s disease (CD) ileum. **B.** Target cells expressing CSF2RA (left) and CSF2RB (right). **C.** Representative Xenium spatial transcriptomic images showing the localization of CSF ligands and receptors in paired uninvolved and inflamed Crohn's disease ileal tissue. Top panels: CSF1 (green), CSF2/GM-CSF (red), and CSF3 (yellow) transcript distribution. Bottom panels: CSF1R (green), CSF2RA (red), and CSF2RB (orange) transcript distribution. Scale bar, 500 μm.

**A**

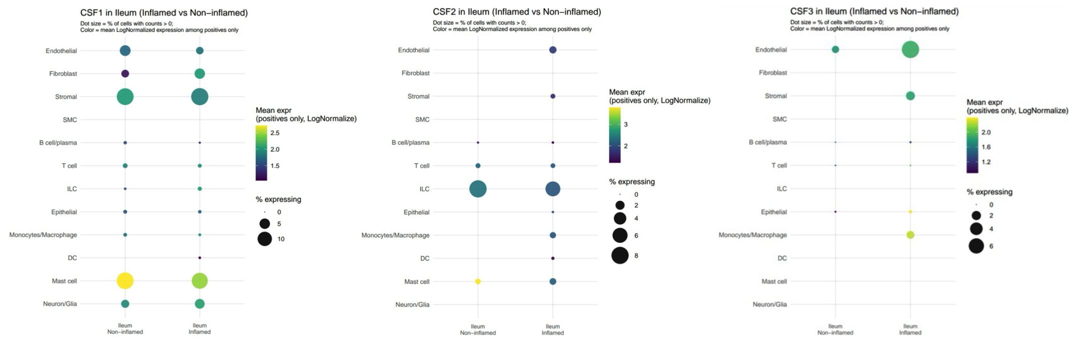


**B**

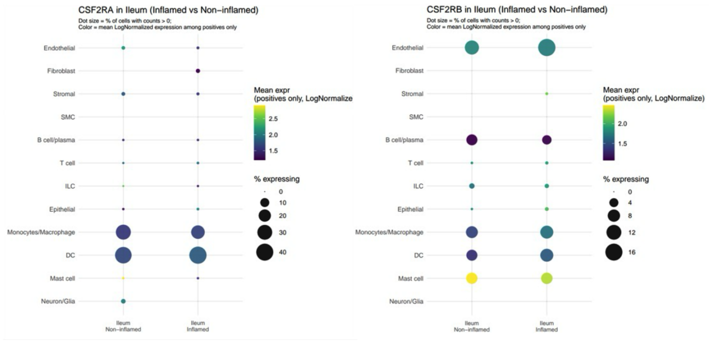


**C.**

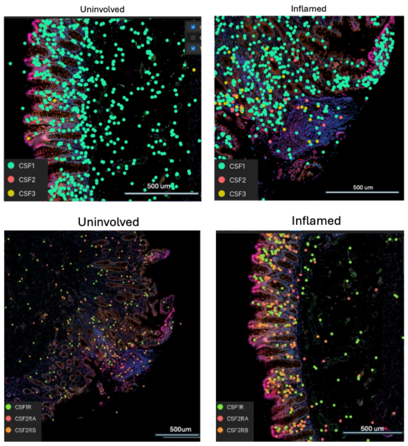

**Supplementary figure 3.** Knockout of zebrafish *csf2rb* using CRISPR-CAS9. **A.** Images of embryos with fluorescently-labeled trans-activating CRISPR RNA. **B.** Human CAS9 protein and fluorescent-labeled gRNA (Atto488 tracrRNA) were co-microinjected into one-cell zebrafish embryos (N=600). The genomic DNA of atto488 tracrRNA positive embryos were collected for T7E1 assay to confirm CAS9 cutting. The F0 Mosaics were incrossed for transmitting germline mutations. F1 were genotypes and the heterozygous mutants were incrossed for homozygous mutant carriers. The mutation was identified as a single nucleotide (G) insertion in exon three and leads to a frameshift and early termination of Csf2rb. **C.** Full DNA gel image of Step 3 CRISPR/Cas9 digestion shown in Figure S3B. The red rectangle highlights the area of interest.

A

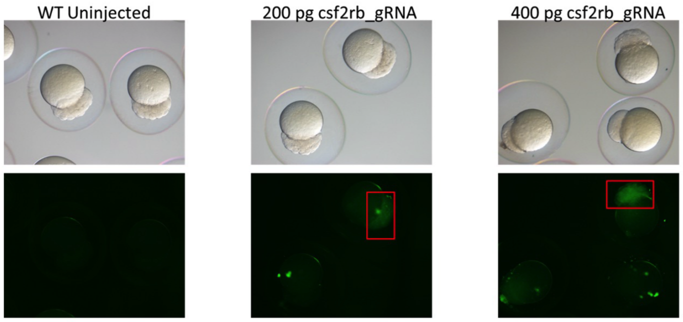


B

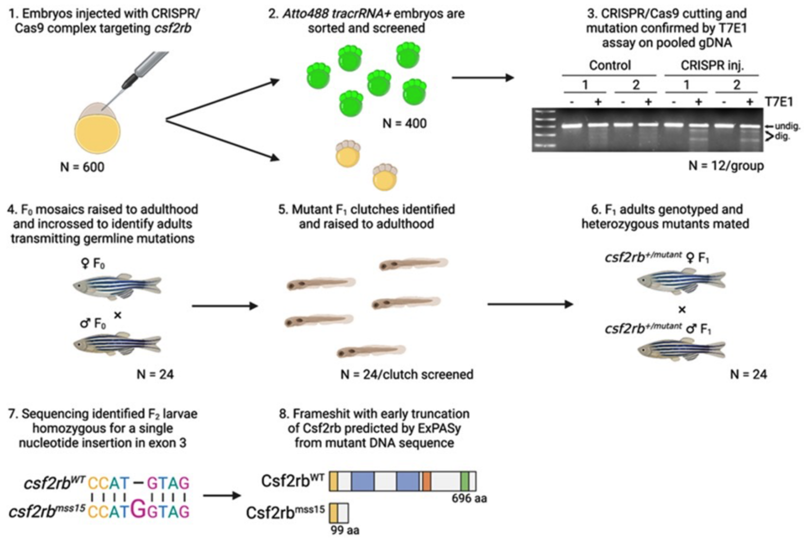


C

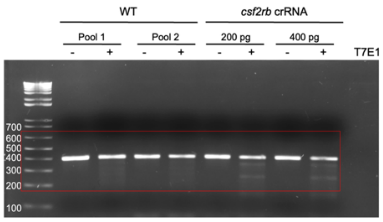

**Supplementary figure 4. A.** RNAscope in situ hybridization of RORC, TNFa, IL1B and OSM in wildtype and CSF2rb-/- zebrafish intestine. Scale bar 50 μm. **B.** No probe control image for csf2rb. **C.** RNAscope in situ hybridization. The quantification of the ILC marker il7r. **D.** No probe control image for il7r/il22/ifng1.1.

**A**

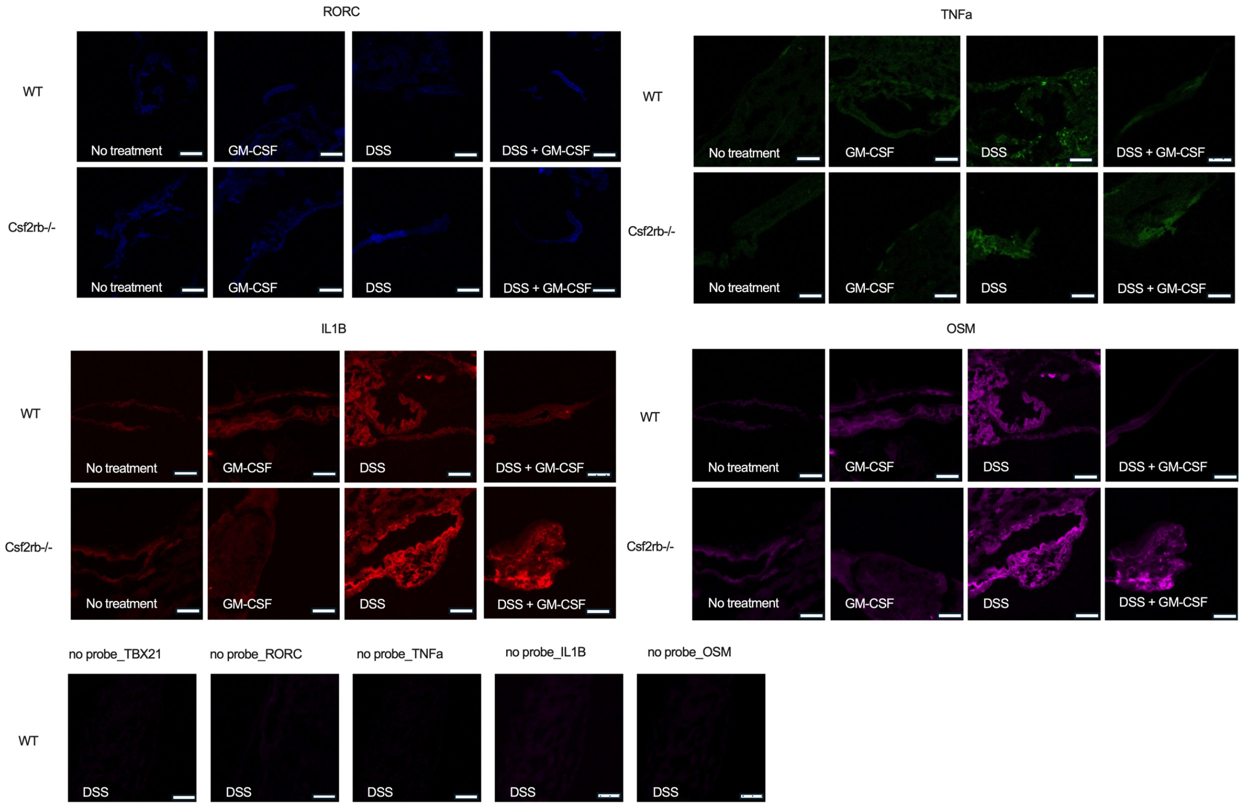


**B**

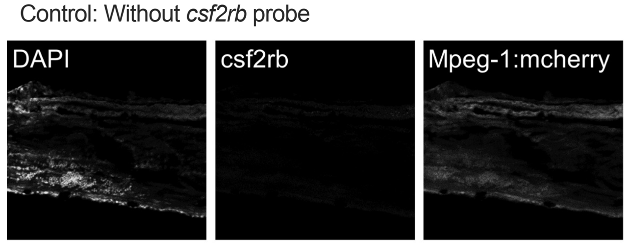


**C**

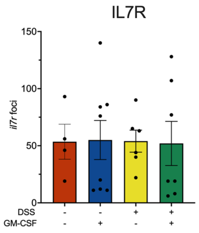


**D**

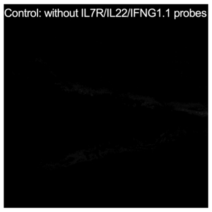

**Supplementary figure 5. A.** Raw spilced/unspliced counts and ratio of top proliferative ILC marker genes. **B.** Xenium image showing identified ILC1s (IL7R+, KLRB1+, TBX21+) and ILC3s (IL7R+, KLRB1+, RORC+), differentiated from T cells and NK cells.

**A**

| Gene | Condition | Spliced raw | Unspliced raw | Total count | Spliced ratio | Unspliced ratio |
|---|---|---|---|---|---|---|
| pfn1 | Control | 30 | 55 | 85 | 35.29 | 64.71 |
| | GM-CSF | 29 | 23 | 52 | 55.77 | 44.23 |
| | DSS | 23 | 20 | 43 | 53.49 | 46.51 |
| | DSS+GM-CSF | 9 | 36 | 45 | 20.00 | 80.00 |
| hmgn2 | Control | 12 | 73 | 85 | 14.12 | 85.88 |
| | GM-CSF | 19 | 30 | 49 | 38.78 | 61.22 |
| | DSS | 38 | 0 | 38 | 100.00 | 0.00 |
| | DSS+GM-CSF | 15 | 29 | 44 | 34.09 | 65.91 |
| rsp17 | Control | 53 | 73 | 126 | 42.06 | 57.94 |
| | GM-CSF | 41 | 18 | 59 | 69.49 | 30.51 |
| | DSS | 42 | 0 | 42 | 100.00 | 0.00 |
| | DSS+GM-CSF | 31 | 15 | 46 | 67.39 | 32.61 |
| arpc1b | Control | 3 | 87 | 90 | 3.33 | 96.67 |
| | GM-CSF | 3 | 52 | 55 | 5.45 | 94.55 |
| | DSS | 37 | 3 | 40 | 92.50 | 7.50 |
| | DSS+GM-CSF | 14 | 26 | 40 | 35.00 | 65.00 |
| h2az2b | Control | 29 | 42 | 71 | 40.85 | 59.15 |
| | GM-CSF | 25 | 28 | 53 | 47.17 | 52.83 |
| | DSS | 41 | 0 | 41 | 100.00 | 0.00 |
| | DSS+GM-CSF | 21 | 15 | 36 | 58.33 | 41.67 |
| hmgb2b | Control | 33 | 28 | 61 | 54.10 | 45.90 |
| | GM-CSF | 22 | 27 | 49 | 44.90 | 55.10 |
| | DSS | 29 | 11 | 40 | 72.50 | 27.50 |
| | DSS+GM-CSF | 16 | 13 | 29 | 55.17 | 44.83 |
| rpl26 | Control | 17 | 14 | 31 | 54.84 | 45.16 |
| | GM-CSF | 1 | 33 | 34 | 2.94 | 97.06 |
| | DSS | 35 | 0 | 35 | 100.00 | 0.00 |
| | DSS+GM-CSF | 23 | 18 | 41 | 56.10 | 43.90 |

**B**

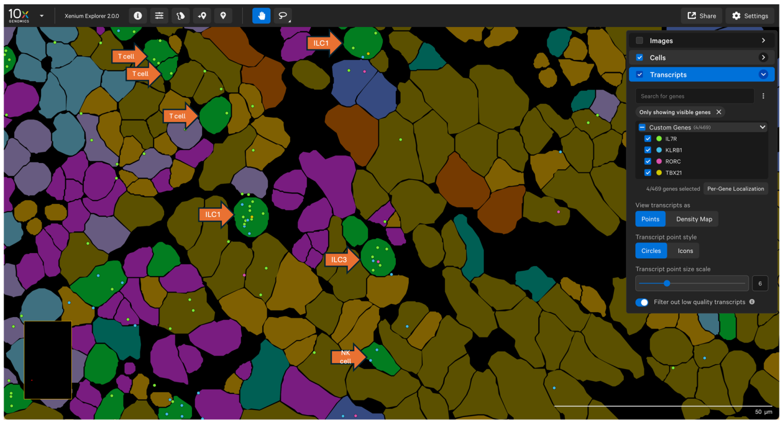

**Supplementary figure 6. A.** H&E staining of inflamed CD tissue sections highlights macrophage aggregates in ulcer exudate (red-circled regions) adjacent to granulation tissue (green-circled regions). **B.** Macrophage aggregates in ulcer exudate were identified through cell clustering and the macrophage marker CD14. **C.** GM-CSF co-localizes with the expression of inflammatory cytokines IL-1β, OSM, and TNF (Arrow) in the inflamed ileum. **D.** No staining control (without primary antibodies) for immunostaining of DAPI, CD14, GM-CSF, pSTAT5, pSTAT3 and pERK1/2. Scale Bar 5000 μm.

**A**

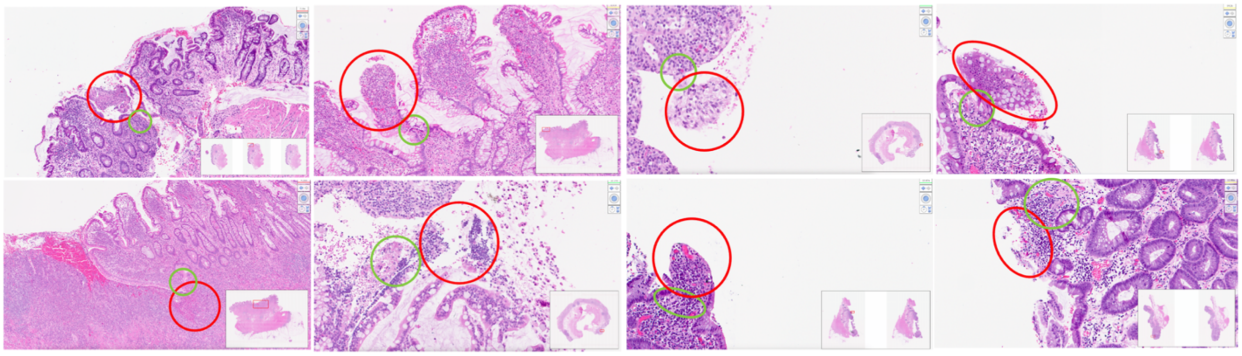

**B**

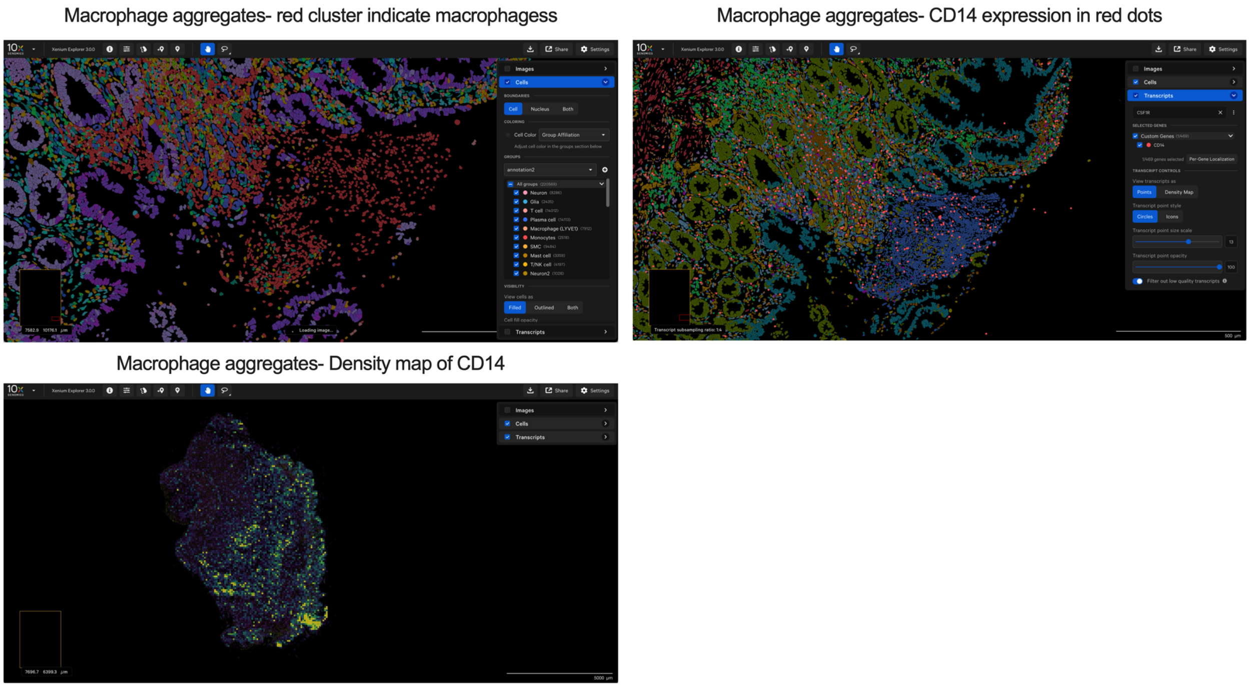

**C**

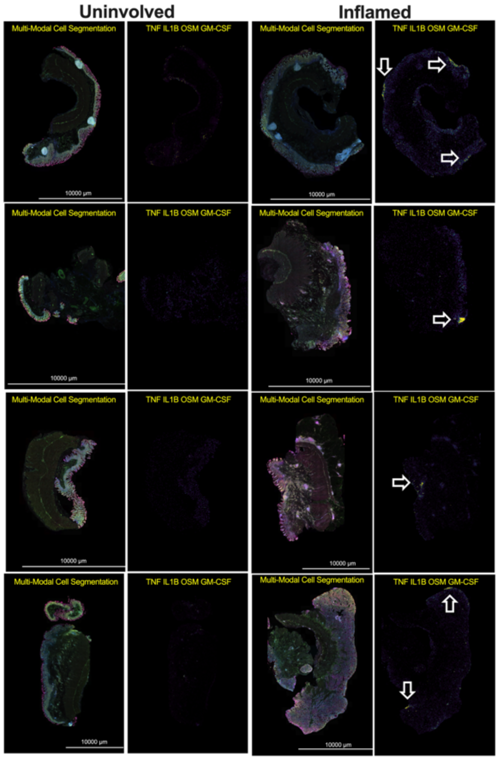


**D**

Control: adjecent section without antibodies

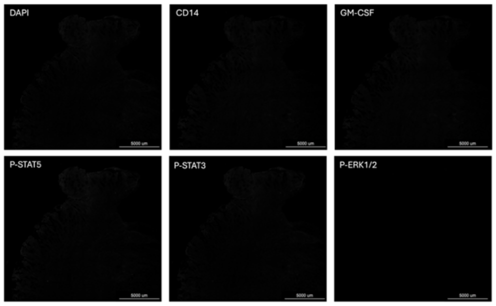

**Supplementary Figure 7. Conserved macrophage transcriptional programs induced by GM-CSF signaling.** Heatmaps summarizing selected macrophage-associated genes across three complementary analyses: (left: GM-CSF response) zebrafish macrophages following DSS + GM-CSF versus DSS alone, (middle: csf2rb dependent) WT versus csf2rb−/− macrophages following DSS and GM-CSF treatment, and (right: Human STAT5-high macrophages) human STAT5-high versus STAT5-low macrophages from Crohn's disease intestine. Genes are grouped according to biological function, including GM-CSF signaling, regulatory/tissue-remodeling macrophage programs, and macrophage pathways associated with establishment of an ILC3-supportive tissue niche. Gray boxes indicate genes that were not detected in the corresponding dataset.

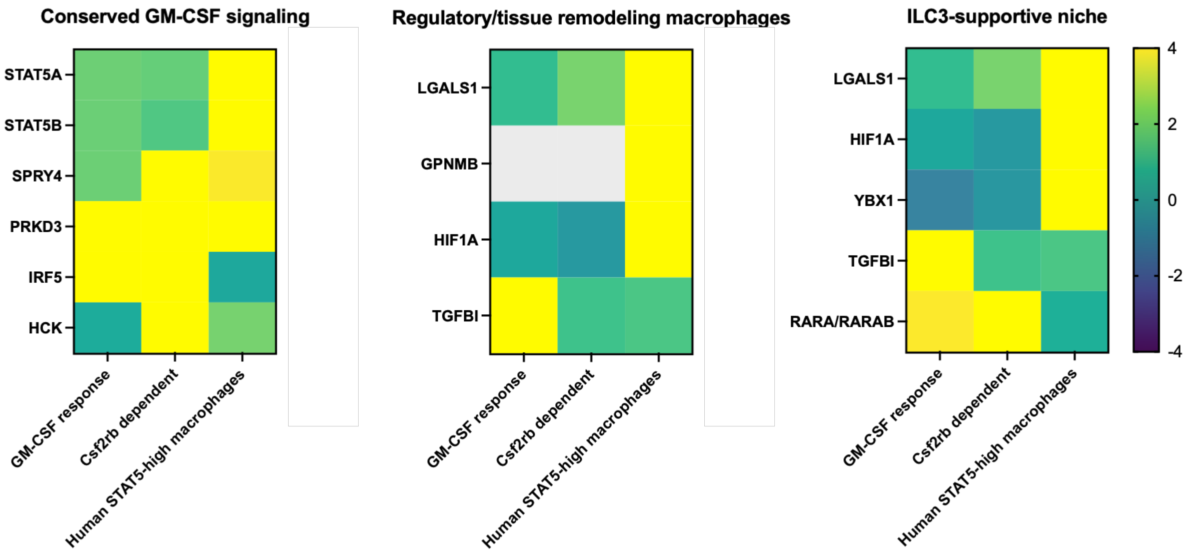

Supplementary table 1. Patient information.

| GRID | BRP ID | Year of diagnosis | Date Collected/Surgery | DOB | Age | Race | Hispanic/Latino | Jewish | Gender | Slide A | Slide B | Slide C | Areas of Sample Collected | Disease Severity | Medications |
|---|---|---|---|---|---|---|---|---|---|---|---|---|---|---|---|
| 1011328 | 50336 | 2022 | 11/27/23 | 01/26/00 | 23 | White | No | Yes | M | Stricture | Involved | Uninvolved | Affected/ Unaffected/ Strictures | B3 | Corticoids; Atb; Anti-TNF |
| 1011332 | 50403 | 1978 | 11/30/23 | 04/29/59 | 64 | White | No | No | F | Uninvolved | Involved | Stricture | Affected/ Unaffected/ Strictures | B2 | Corticoids; Atb; Anti-TNF |
| 1011325 | 50217 | 1993 | 11/17/23 | 04/16/76 | 47 | White | No | Yes | F | Involved | Uninvolved | Stricture | Affected/ Unaffected/ Strictures | B3 | ASA; Corticoids; Atb; Immunomodulatory drugs; Anti-TNF |
| 1011312 | 50006 | 2015 | 11/8/23 | 09/16/48 | 75 | White | No | No | F | Involved | Uninvolved | Stricture | Affected/Unaffected/ Strictures | B2 | Corticoids; Atb; Immunomodulatory drugs; MTX |
| 1011335 | 50452 | 1970 | 12/4/23 | 3/15/54 | 69 | White | No | No | F | Uninvolved | Involved | Stricture | Affected/Unaffected/Strictures | B2 | Corticosteroids,Aminosalycates, Abx |
| 1011431 | 52070 | 2003 | 3/4/24 | 2/14/79 | 46 | White | No | No | F | Involved | Uninvolved | Stricture | Affected/ Unaffected/ Strictures | B1 | Oral Steroids, Immunomodulatory drugs ,Biologicsand Anti-TNF |
| 1011455 | 52583 | 2015 | 3/25/24 | 02/10/00 | 25 | White | No | Yes | M | Involved | Uninvolved | Stricture | Affected/Unaffected/Strictures | B2 | Oral Steroids, Immunomodulatory drugs and Anti-TNF |
| 1011354 | 50607 | 1991 | 12/13/23 | 03/31/72 | 53 | White | No | No | F | Uninvolved | Involved | Stricture | Affected/Unaffected/Strictures | B2 | Oral Steroids and Immunomodulatory drugs |
| 1011369 | 51069 | 1993 | 1/11/24 | 03/29/59 | 66 | White | No | No | F | Uninvolved | Involved | Stricture | Affected/Unaffected/Strictures | B2 | Oral Steroids |
| 1011564 | 53338 | 1987 | 4/26/24 | 02/04/56 | 69 | White | No | No | M | Involved | Uninvolved | Stricture | Affected/Unaffected/Strictures | B1 | ASA, Immunomods, Anti-TNF,Other Biologics |
